# Transcriptional Circuit Fragility Influences HIV Proviral Fate

**DOI:** 10.1101/504969

**Authors:** Emily L. Morton, Christian V. Forst, Yue Zheng, Ana B. De Paula-Silva, Nora-Guadalupe P. Ramirez, Vicente Planelles, Iván D’Orso

**Affiliations:** Department of Microbiology, University of Texas Southwestern Medical Center, Dallas, TX 75390; Department of Genetics and Genomic Sciences, Institute for Genomics and Multiscale Biology, Icahn School of Medicine at Mount Sinai, New York, NY 10029; Department of Pathology, University of Utah, Salt Lake City, UT 84112; Peloton Therapeutics, 2330 Inwood Rd #226, Dallas, TX 75235

## Abstract

Transcriptional circuit architectures can be evolutionarily selected to precisely dictate a given response. Unlike these cellular systems, HIV is regulated through a complex circuit composed of two successive phases (host and viral), which create a positive feedback loop facilitating viral replication. However, it has long remained unclear whether both phases operate identically and to what extent the host phase influences the entire circuit. Here we report that while the host phase is regulated by a checkpoint whereby KAP1 mediates transcription activation, the virus evolved a minimalist system bypassing KAP1. Given the complex circuit’s architecture, cell-to-cell KAP1 fluctuations impart heterogeneity in the host transcriptional responses thus affecting the feedback loop. Mathematical modeling of a complete circuit reveals how these oscillations ultimately influence homogeneous reactivation potential of a latent virus. Thus, while HIV drives molecular innovation to fuel robust gene activation, it experiences transcriptional fragility thereby influencing viral fate and cure efforts.

**In Brief:** HIV evolved a minimalist but robust transcriptional circuit bypassing host regulatory checkpoints; however, the fragility of the circuit in the host phase (which primes HIV for activation) largely affects proviral transcription and fate.

**Highlights:**

- The host and viral phases of the HIV transcriptional circuit have different functional requirements
- HIV evolved a minimalist program to robustly bypass host cell regulatory checkpoints
- A mathematical model reveals that the host phase is subject to transcriptional circuit fragility
- Host transcriptional circuit fragility influences the viral feedback and latency reversal potential

## INTRODUCTION

Transcriptional regulatory circuits are essential for several key biological processes such as development, differentiation and cell fate responses. As such, transcriptional circuit architecture can be evolutionarily selected to precisely dictate a given response. In contrast to these highly evolvable circuits, viruses like human immunodeficiency virus type-1 (HIV), which integrate into the human genome (Hughes and Coffin, 2016; Schroder et al., 2002), initially fall under the control of host circuits. Given HIV integration is “quasi”-random, the heterogeneous integration landscape may impact transcriptional circuit architecture leading to variable outcomes (activation or silencing) thereby generating profound phenotypic diversity among different infections (i.e., active or latent), here referred to as “proviral fate” (**Figure 1A**).

**Figure 1.**
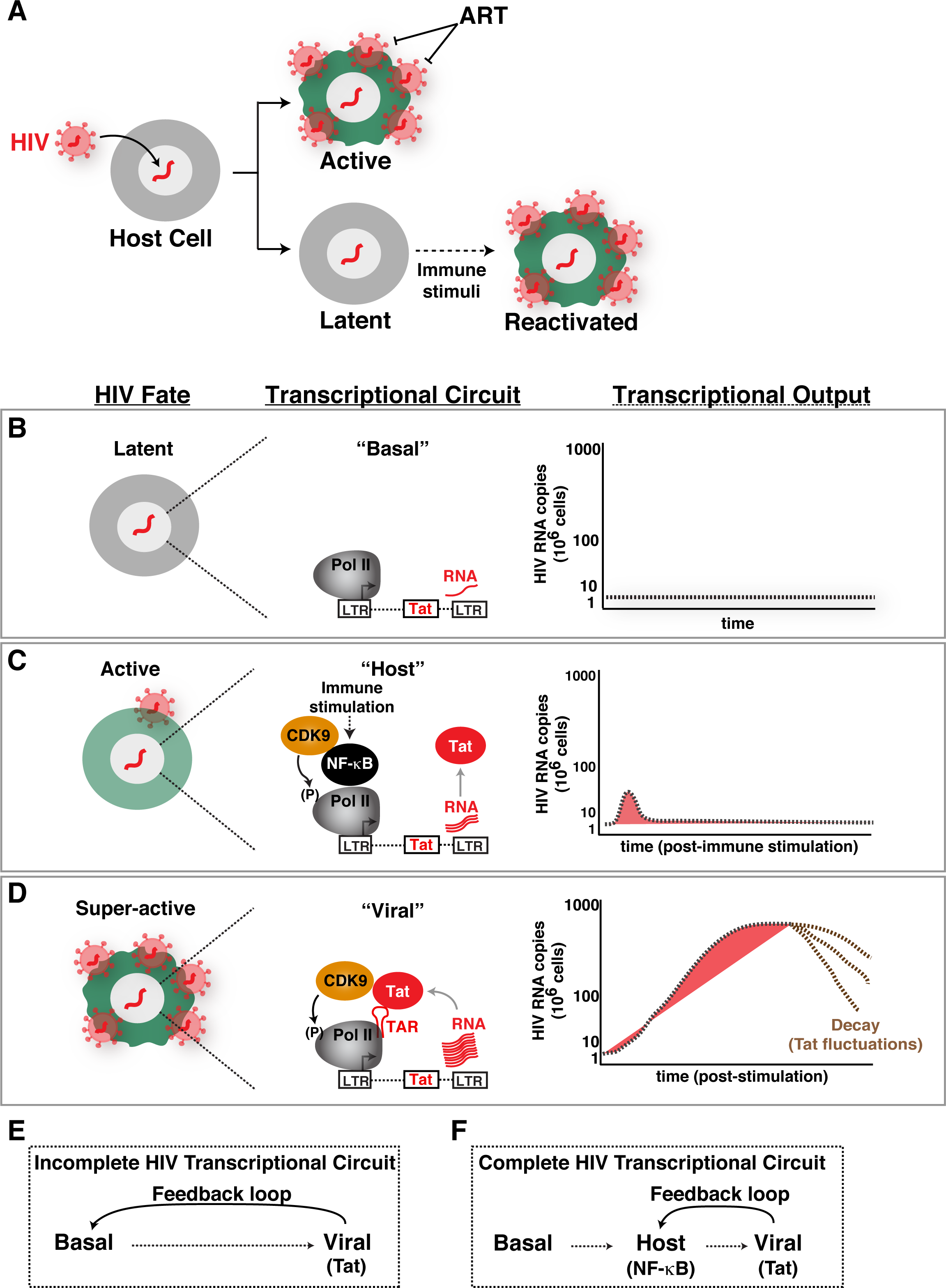
Establishing an Experimental – Mathematical Modeling Framework for Understanding a Complete HIV Transcriptional Circuit. (A) Simplistic scheme of HIV proviral fate (active or latent) after infection and integration into the host cell genome. Latent viruses can be reactivated in response to immune stimulation. (B–D) Schemes depicting three different proviral fates (latent, active, super-active), and their associated transcriptional circuits (basal, host, viral) and outputs. See text for complete details. (E) Scheme of an incomplete HIV transcriptional circuit as previously interrogated by Weinberger et al. (Weinberger et al., 2005). (F) Scheme of a complete HIV transcriptional circuit as interrogated in this work.

Over the past three decades, one of the most exciting breakthroughs in biomedical research was the discovery of anti-retroviral therapy (ART), which suppresses active viral replication to nearly undetectable levels. However, ART fails to cure latent infections because the targeted proteins are not expressed or expressed at extremely low levels. Consequently, HIV establishes long-lived latent reservoirs *in vivo* by persisting as a stable integrated provirus in resting memory CD4+ T lymphocytes and myeloid cells (Eisele and Siliciano, 2012; Margolis, 2010; Pierson et al., 2000), and by remaining undetected by immune surveillance mechanisms. Although these constitute a very small population (∼1 in a million CD4+ cells), they do not apparently produce appreciable virus, and are considered the largest barrier for HIV eradication from a patient (Chun et al., 1997; Chun et al., 1995; Finzi et al., 1999).

Although the molecular rules governing proviral latency appear to be pleiotropic (Dahabieh et al., 2015; Lassen et al., 2004; Ruelas and Greene, 2013), one common feature is the resting state of the infected cell leading to low, or even undetectable, levels of transcription activity. Thus, HIV latency is a state of non-productive infection due to restrictions, primarily, at the transcriptional level (Dahabieh et al., 2015; Karn, 2011; Lassen et al., 2004; Margolis, 2010; Ruelas and Greene, 2013; Siliciano and Greene, 2011).

Because cessation of therapy leads to virus rebound within weeks, HIV infected individuals must remain on therapy permanently (Eisele and Siliciano, 2012; Siliciano and Greene, 2011). Given the secondary effects associated with the long-term regime (including toxicities, aging, and other associated diseases), pharmacological strategies designed to eradicate the viral latent reservoir represent a critical unmet need. There is enormous enthusiasm for the potential of precision therapies targeting the latent reservoir in clinical settings. Thus, HIV latency has become the center of attention. As such, a large body of research has identified the role of individual host factors and epigenetics on HIV transcription activation or silencing, and elucidated host cell enzymes as targets that could be manipulated using chemical probes to induce reactivation of latent viruses. Despite several landmark discoveries, we currently lack a complete understanding of the fundamental regulatory principles of the HIV transcriptional circuit and its implications for proviral fate control, including latency.

The HIV transcriptional circuit is regulated at different levels. First, during normal cell homeostasis, “basal” steady-state transcription maintains a low-level of non-productive RNA synthesis leading to short, immature transcripts (**Figure 1B**). In this state, the viral activator Tat is not expressed and, thus, HIV does not replicate (latent state). In the “host” phase, when cells are exposed to immune stimulation, transcription factors like NF-κB and NFAT are activated leading to an initial, low-level “boost” in proviral transcription. In proviruses lacking Tat, this phase shows a unimodal pattern of activation that is quickly turned-off leading to a small amount of viral products (Feinberg et al., 1991) (**Figure 1C**). During productive infections with wild-type proviruses, the initial transcriptional boost is critical because it enables Tat synthesis before the host phase turns off. In this case, the host phase is rapidly followed by a “viral” phase in which Tat amplifies transcription by more than 100-fold promoting a positive transcriptional feedback loop and robust viral replication (Feinberg et al., 1991; Jordan et al., 2001; Kao et al., 1987; Karn, 1999, 2011; Laspia et al., 1989; Ott et al., 2011; Rice and Mathews, 1988) (**Figure 1D**). It is believed that under special circumstances where the cell state changes (e.g., cellular relaxation inducing quiescence) Tat fluctuations lead to transcriptional decay thereby promoting latency establishment (Weinberger et al., 2005).

In the resting scenario, most of the cellular activators are found in a latent state but they become activated when the infected immune cells encounter a stimulus from the microenvironment. For example, the pro-inflammatory cytokine <u>T</u>umor <u>N</u>ecrosis <u>F</u>actor (TNF) activates the canonical NF-κB pathway leading to phosphorylation of the RelA subunit (hereafter referred to as NF-κB), and translocation from the cytoplasm into the nucleus where it recognizes its binding element at the viral promoter driving proviral transcription (Nabel and Baltimore, 1987; Williams et al., 2006; Williams et al., 2007). Similarly, the CD40 ligand and lymphotoxin (LTα and LTβ) induce proviral transcription upon receptor activation and signaling through the non-canonical NF-κB pathway (Pache et al., 2015). T-cell stimulation functions broadly through multiple signaling pathways including the cellular activators NF-κB, NFAT, and AP-1 (Duverger et al., 2013; Kinoshita et al., 1997; Nabel and Baltimore, 1987).

Both cellular activators and Tat promote proviral transcription in the host and viral phases, respectively, through a complex layer of host factors including general transcription factors, RNA polymerase II (Pol II), co-activators/co-repressors, and chromatin-modifying enzymes, among others (Ott et al., 2011; Van Lint et al., 2013). One key host factor is the positive transcription elongation factor b (the P-TEFb kinase) (Dalal and Johnson, 2017; Wei et al., 1998), which is composed of a cyclin T subunit and the catalytic CDK9 subunit (hereafter referred as CDK9). Both cellular activators and Tat utilize CDK9 to facilitate the transcription elongation program, a critical step in the viral life cycle (Mancebo et al., 1997; Ott et al., 2011; Peterlin and Price, 2006; Zhou et al., 2000). In contrast to the cellular activators that bind *cis*-elements at the viral promoter, Tat binds the TAR (<u>T</u>rans<u>a</u>ctivation <u>R</u>esponse element) RNA structure formed on nascent viral pre-mRNAs directly recruiting P-TEFb to phosphorylate Pol II and the negative elongation factors to promote productive elongation (Feinberg et al., 1991; Mancebo et al., 1997; McNamara et al., 2016a; Ott et al., 2011).

Despite the relevance of CDK9 for activation of, both, host and viral phases of the HIV transcriptional circuit, it has long remained unclear whether both programs operate through identical or different mechanisms and how their malfunction impact proviral latency. Interestingly, recent studies have suggested a role for the transcriptional regulator KAP1 (also known as TRIM28, TIF1β) in proviral transcription through CDK9 recruitment to the promoter as part of the 7SK complex, in which the kinase remains in a primed state (D’Orso, 2016; McNamara et al., 2016a; McNamara et al., 2016b; Michels et al., 2004). In this context, the 7SK complex (composed of 7SK RNA and kinase inhibitor HEXIM) not only inactivates the kinase, but more importantly, it has a positive role in delivering the kinase for “on site” activation at the viral promoter (McNamara et al., 2016b). These recent discoveries provide an unprecedented function for KAP1, which has been previously implicated in transcriptional repression through epigenetic silencing of genes and retroelements in progenitor and non-committed cells as well as repression of viruses in embryonic stem (ES) cells (Brattas et al., 2017; Ellis et al., 2007; Rowe et al., 2013; Wolf and Goff, 2007; Wolf et al., 2008).

Notably, here we found that KAP1 is expressed in primary resting memory CD4+ T cells (one of the largest reservoirs of latent HIV in infected individuals), and is recruited to the proviral genome in this primary cell model, thus providing biological relevance for the pathogenic mechanism herein described. To our surprise, we also report the unexpected findings that the different phases of the HIV circuit have different functional requirements. While KAP1 is critical for activation of the host phase, HIV evolved a minimalist system where Tat represents a switch to a ‘higher gear’ bypassing KAP1 to activate proviral transcription. While KAP1 recruits CDK9 to the promoter to facilitate activation by cellular activators in response to cytokine stimulation, Tat subsequently functions in a KAP1-independent manner directly recruiting the kinase to sustain transcription elongation. Given the host phase has a strict requirement for KAP1, its loss directly impacts on the positive feedback loop, thus reducing the magnitude of reactivation of a latent virus.

Previous studies have created mathematical models that incompletely interrogate the HIV transcriptional circuit (“basal-viral”) (Weinberger et al., 2005) (**Figure 1E**). Thus, the roles of host cell factors and immune cell stimulation on the host phase and its effect on the positive feedback loop have not been previously interrogated. Given the virus strictly relies on the immune cells’ activation status, we rationalized that generating a model that can recapitulate the complete HIV program cannot only provide critical insights into HIV biology, but also pave the groundwork for more efficient interventions in the clinical setting. We thus created the first mathematical model that recapitulates the complete HIV transcriptional circuit (“basal-host-viral”). This model predicts that fluctuations of KAP1 levels in patient’s cells could affect the host phase; and, as a consequence, the magnitude of the Tat feedback (thereby dampening latency reversal potential). We tested this model in several clonal cell lines and observed how KAP1 protein oscillations impart heterogeneity in the transcriptional responses thereby influencing the reactivation potential of a latent virus. Thus, cell-to-cell fluctuations in the levels of key components of the transcription machinery in patients under suppressive therapy could affect “homogeneous” reactivation potential. Our findings provide a mechanistic explanation for the importance of the host phase to ensure the virus is readily and robustly activated during infection to complete the pathogenic cycle. While HIV drives molecular innovation to fuel robust gene activation, it suffers from host transcriptional circuit fragility thereby influencing proviral transcription and fate.

## RESULTS

### Establishing an Experimental – Mathematical Modeling Framework for Interrogating a Complete HIV Transcriptional Circuit

HIV infection of target immune cells can lead to active and latent infections as a potential consequence of transcriptionally active and silent states, respectively (**Figure 1A**). **Figures 1B–D** illustrate the progression of molecular events leading to activation of the HIV transcriptional circuit from basal transcription, to activation of the host phase by cellular activators (e.g., NF-κB) in response to immune stimulation; and ultimately, to activation of the viral phase by Tat. The key feature of this system is that activation of the host phase during productive infection leads to Tat synthesis, which induces a positive feedback loop (Tat feedback) leading to robust viral products synthesis and replication. The activators of both phases (NF-κB and Tat) are believed to function by recruiting the transcription elongation complex (CDK9) to the integrated proviral genome to induce Pol II release from the pause site at the promoter; and consequently, productive transcription elongation.

Because the HIV circuit operates through the combined, sequential activity of the host and viral phases, it has been challenging to uncouple the precise contributions of each phase in the program as a whole. To overcome this challenge, here we establish an integrated experimental and mathematical modeling framework for precisely interrogating a complete HIV transcriptional circuit. Experimentally, we use both primary cells and transformed cell-based models containing integrated HIV where Tat can be either wild-type (Tat+) or defective (Tat-), which are regulated by the complete circuit (“host” and “viral” phases) or by the minimalist version of it (“host” phase only), respectively. Directly comparing the transcriptional profiles of both proviruses in response to immune stimulation upon host cell factor depletion enable us to infer their contributions to the different phases of the HIV transcriptional program.

To expand the establishment of experimental approaches, a mathematical modeling was created that recapitulates, for the first time, the complete architecture (“basal-host-viral”) of the HIV transcriptional circuit (**Figure 1F**). While previous studies have modeled HIV activation by Tat using a simple circuit composed of the “basal-viral” phases (**Figure 1E**) (Weinberger et al., 2005; Weinberger et al., 2008), those models do not enable one to examine the contribution of the host phase to the viral phase, the magnitude of the feedback loop nor the reactivation potential of a latent virus. Given this large caveat, it has been impossible to predict and test what are the contributions of host cell factors to the HIV transcriptional program as a whole.

Taken together, our combined experimental and computational framework establishes unique and tractable models to interrogate the complete HIV circuit and its implications in the context of proviral latency and reactivation. Here, we examine the contributions of host cell factors implicated in proviral transcriptional elongation control to various HIV circuit architectures.

### Defining Host Cell Factor Contributions to the Transcriptional Circuit of a Latent Virus

KAP1 has been previously shown to play an important role in epigenetic silencing of retroelements and genes in ES cells and progenitor cells (Brattas et al., 2017; Ellis et al., 2007; Rowe et al., 2010; Rowe et al., 2013; Wolf and Goff, 2007, 2008, 2009). As part of this mechanism, KAP1 appears to be recruited to chromatin (promoter regions) through interaction with a family of KRAB-domain zinc finger proteins (KRAB-ZnF) (Iyengar and Farnham, 2011). Then, KAP1 recruits chromatin-modifying enzymes that promote epigenetic silencing, such as SETDB1 (H3K9 tri-methylase) and the NURD/HDAC complex (known to promote histone deacetylation) to ultimately silence the locus (**Figure 2A**) (Iyengar and Farnham, 2011; Karimi et al., 2011; Macfarlan et al., 2011; Matsui et al., 2010).

**Figure 2.**
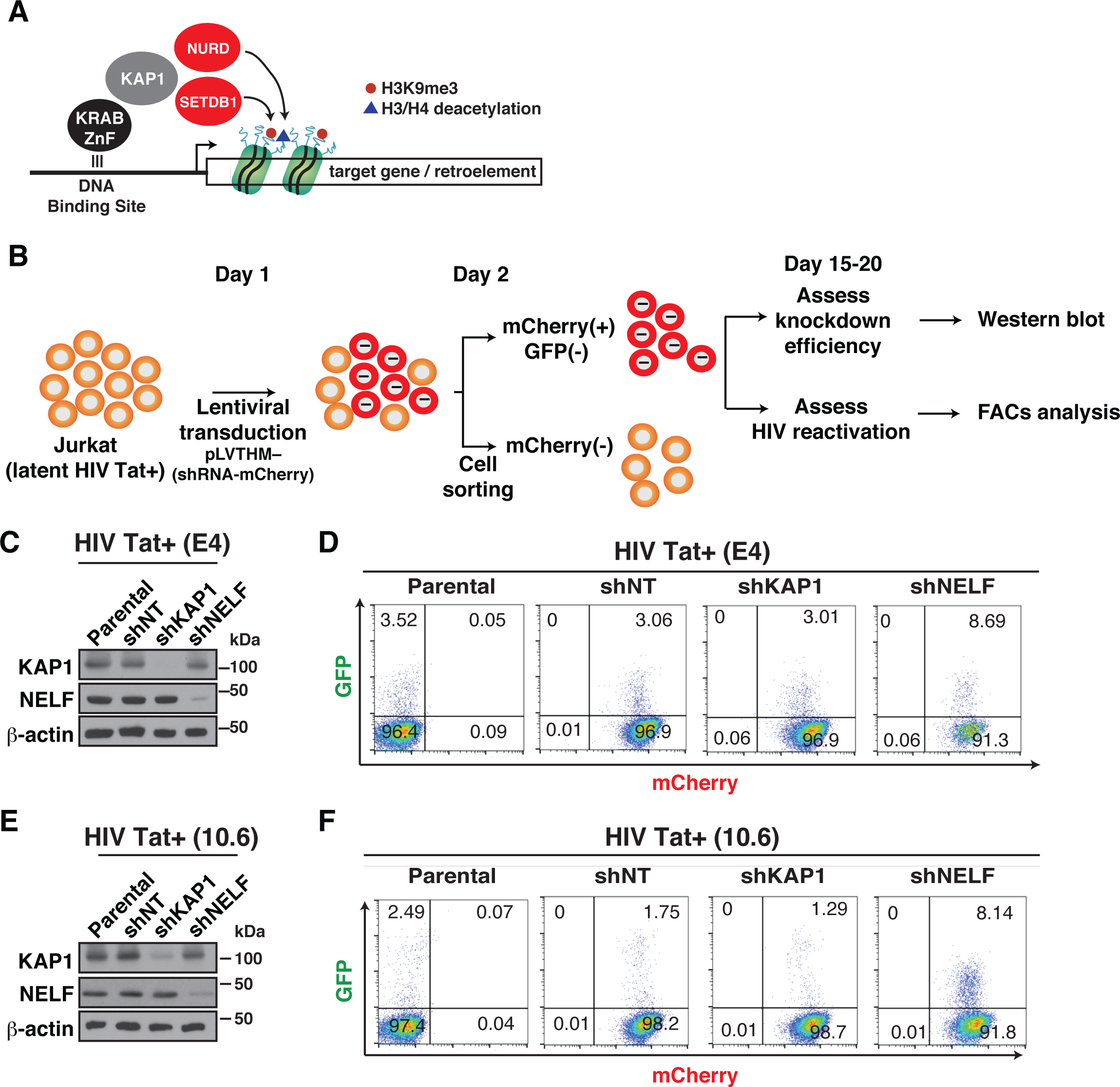
Loss of KAP1 Does not Reactivate HIV from Cell-based Models of Latency. (A) Simplified current model of KAP1-mediated transcriptional silencing based on data from several studies (Iyengar and Farnham, 2011; Karimi et al., 2011; Macfarlan et al., 2011; Matsui et al., 2010; Rowe et al., 2010; Rowe et al., 2013). (B) Overview of the protocol used to transduce and analyze by cell sorting the cell-based models of HIV latency using lentiviral vectors co-expressing shRNA and mCherry. (C) Western blots of the HIV Tat+ (E4) cell-based models with the indicated antibodies. (D) Flow cytometry analysis (GFP and mCherry) of the HIV Tat+ cell-based models from panel (C). The number of GFP+ cells from three independent runs were: 3.52 ± 0.61 (Parental), 3.06 ± 0.43 (shNT), 3.01 ± 0.37 (shKAP1), and 8.69 ± 1.08 (shNELF). (E) Western blots of the HIV Tat+ (10.6) cell-based models with the indicated antibodies. (F) Flow cytometry analysis (GFP and mCherry) of the HIV Tat+ cell-based models from panel (E). The number of GFP+ cells from three independent runs were: 2.49 ± 0.66 (Parental), 1.75 ± 0.11 (shNT), 1.29 ± 0.07 (shKAP1), and 8.14 ± 0.87 (shNELF).

To test the hypothesis that KAP1 is recruited to the HIV 5’-long terminal repeat (LTR) to promote proviral latency through epigenetic silencing of the host phase of the circuit, we transduced the Jurkat cell-based models of latency E4 and 10.6, which contain Tat (HIV Tat+ provirus) and a GFP marker for ease of measurement (Jordan et al., 2003; Pearson et al., 2008), with self-inactivating lentiviruses (pLVTHM) expressing non-targeting (NT) or KAP1 shRNAs (**Figure 2B**). As a positive control for our experiments, we used a shRNA targeting the NELF-E subunit of the Negative Elongation Factor Complex (NELF), which has been shown to relieve Pol II pausing at the HIV promoter to spontaneously induce proviral transcription and latency-reversal (Jadlowsky et al., 2014; Kaczmarek Michaels et al., 2015). Given the pLVTHM vector co-expresses the shRNA and a fluorescent marker (mCherry) we used <u>F</u>luorescent <u>A</u>ctivated <u>C</u>ell <u>S</u>orting (FACS) to separate efficiently transduced cells (mCherry+ and GFP-) from untransduced cells (mCherry- and GFP-) to assess knockdown (KD) efficiency in the population by western bot analysis, and reactivation of latent HIV by single-cell flow cytometry analysis by computing the number of GFP+ cells (**Figures 2C–F**).

Despite the remarkably efficient KD of KAP1 in the HIV Tat+ (E4) cell-based model (>90% KD compared to the shNT cell line), we did not observe reactivation of latent HIV as revealed by the similar levels of GFP+ cells in the parental, NT, and KAP1 shRNA cell lines (**Figures 2C and 2D**). However, as expected, loss of NELF resulted in an increase in the percentage of GFP+ cells (∼2.8-fold increase over NT; **Figure 2D**), consistent with previous studies (Jadlowsky et al., 2014; Kaczmarek Michaels et al., 2015).

While the previous analysis was performed in a bulk population, we sought to determine whether these results could be reproduced at a single-cell level. To that end, we sorted individual cells, generated clonal cell lines and examined by FACS their latency-reversal potential. Consistent with the results obtained at the population level, individual shKAP1 clones did not show latent HIV reactivation compared to shNT clones (data not shown). In addition, we found great variability in the reactivation potential in NELF-depleted clones (data not shown), as previously reported by the Karn lab (Jadlowsky et al., 2014), potentially implying heterogeneity in clonal responses due to NELF inactivation. Although understanding the mechanisms would be interesting, this is beyond the scope of this study.

To test if the results obtained in the HIV Tat+ (E4) system were cell model independent and thus generalizable, we recreated a collection of cell lines on the 10.6 cell-based model generated by the Verdin lab (Jordan et al., 2003). Consistent with the data obtained in E4, we did not observe significant changes in the percentage of GFP+ cells between parental, shNT and shKAP1 10.6 cell lines (**Figures 2E and 2F**). Again, efficient NELF KD in the 10.6 cell-based model (shNELF) led to an increase in the number of GFP+ cells compared to both parental and shNT cell lines (**Figure 2F**), indicating reactivation of latent HIV, with slightly higher reactivation levels in 10.6 cells compared with those in E4 cells (∼4.7-fold vs ∼2.8-fold, respectively), probably due to intrinsic differences in the two systems (**Figures 2D and 2F**).

Given HIV integrates semi-randomly and can be found in sites with different chromatin accessibility (Hughes and Coffin, 2016; Jordan et al., 2003; Jordan et al., 2001; Maldarelli et al., 2014; Schroder et al., 2002), we asked whether KAP1 contributes to epigenetic silencing in models where the HIV LTR is relatively inaccessible. To test this, we silenced the expression of KAP1 in cell-based models showing lower promoter chromatin accessibility (6.3, 8.4, and 9.2) compared to the previous models (E4 and 10.6) (Jordan et al., 2001; Pearson et al., 2008). Despite the efficient (>95%) loss of KAP1 in every cell-based model examined, as revealed by western blot analysis (**Figure S1A**), we observed no significant changes in the percentage of GFP+ cells in the shKAP1 cell line compared to both parental and shNT cell lines (**Figure S1B**), strongly indicating that KAP1 does not contribute to proviral latency maintenance at least in this permissive CD4+ T cell-based model.

The expression of GFP protein encoded by the proviruses in these cell-based models requires Tat activity and high levels of transcription (Pearson et al., 2008; Weinberger et al., 2005). Thus, it remains possible that loss of KAP1 could promote some degree of latency-reversal but at levels below the GFP detection threshold. To test this possibility, we developed a quantitative real-time PCR (RT-qPCR) assay based on methods that allow us to accurately and efficiently purify and quantitate short (∼17-200 nt), promoter-proximal HIV transcripts (indicative of transcription initiation) and long, promoter-distal HIV transcripts (indicative of transcription elongation), irrespective of the amount of total RNA inputted in the reaction mixture, in the absence/presence of immune stimulation (TNF) (**Figures S1C-F**) (see **Experimental Procedures** for details). The short promoter-proximal transcription start site (TSS)-associated transcripts, are RNA species generated by paused Pol II and potentially held within early elongation complexes (Henriques et al., 2013).

Additionally, to provide further evidence that the amplicons were correctly amplifying the initiating and elongating transcripts we pre-treated HIV Tat+ (E4) cells for 30 min with the potent inhibitor of transcription initiation Triptolide (TRP), which blocks the ATPase activity of the TFIIH helicase XPB, thereby preventing the opening of template DNA (Titov et al., 2011; Vispe et al., 2009), and the inhibitor of transcription elongation Flavopiridol (FP), which blocks the P-TEFb kinase thereby preventing Pol II phosphorylation (Chao and Price, 2001). We observed that TRP, expectedly, blocks TNF-mediated induction of both short (promoter-proximal) and long (promoter-distal) transcripts, while FP only interferes with the synthesis of long (promoter-distal) transcripts, consistent with an initiation and elongation block, respectively (**Figures S1G and H**).

Using this robust, quantitative method, we detected that loss of KAP1 does not promote reactivation of latent HIV in the absence of immune stimulation, consistent with the FACS data. KAP1 KD shows a slight (<1.5-fold), non-significant decrease in the levels of both classes of transcripts in several HIV Tat+ cell-based models (**Figure S1I**), indicating that KAP1 major’s role is not to promote transcription control under basal conditions (McNamara et al., 2016b), but to allow transcriptional responses to immune cell-signaling (see below).

Together, these data suggest KAP1 does not have the expected epigenetic silencing function as seen in other contexts (Brattas et al., 2017; Ellis et al., 2007; Rowe et al., 2010; Rowe et al., 2013; Wolf and Goff, 2007, 2008, 2009). This unexpected result explains why loss of KAP1 does not promote HIV latency reversal as has been seen in other viral systems in non-permissive cells in the absence of immune stimulation (Rauwel et al., 2015); thus, opening the possibility that KAP1 could have other roles in the HIV transcriptional program (e.g., activation) beyond its well-characterized repressor function (see below).

### KAP1 is Expressed and Recruited along with the Host Transcription Elongation Complex to the HIV Genome in Primary Resting CD4+ T Cells

The above data showed KAP1 does not mediate epigenetic silencing of the HIV transcriptional genome in CD4+ T cells as it does with retroelements in ES cells and a subset of target genes during development (e.g., in neuronal progenitor cells) (Brattas et al., 2017; Ellis et al., 2007; Rowe et al., 2010; Rowe et al., 2013; Wolf and Goff, 2007, 2008, 2009). However, our initial experiments were performed in transformed cell-based models and in the absence of immune stimulation, which is key for robust activation of the HIV transcriptional circuit (**Figure 1**). Given this system may not completely recapitulate the establishment and maintenance of latency in primary CD4+ T cells from patients (Chun et al., 1997; Chun et al., 1995; Finzi et al., 1999; Spina et al., 2013), we thus wanted to test whether: (1) KAP1 expression changes in response to T cell state alterations (active vs resting) and HIV infection, (2) KAP1 (and the transcription elongation complex, CDK9) is recruited to the proviral genome in the active and resting T cell states, and (3) KAP1 could drive HIV transcription activation in a more biologically relevant setting in the absence and/or presence of immune stimulation.

To address the first point, we used the primary model in central memory CD4+ T cells (T_CM_) (Bosque and Planelles, 2009; Martins et al., 2016; Spina et al., 2013). Briefly, naïve CD4+ T cells from healthy donors are activated and induced to differentiate into central memory, infected with replication competent virus (HIV-1_NL4.3_) or mock infected (here referred to as “uninfected”), and active infections (p24-positive/CD4-negative) are excluded through magnetic sorting to enrich in latent/uninfected states (**Figures 3A and 3B**). These cells are then maintained in the active T cell state or allowed to transition into a memory resting state in the presence of ART to better mimic viral suppression in patients, and thus create four different experimental groups: (1) “Uninfected (Activated)”, (2) “Productively infected (Activated)”, (3) “Uninfected (Resting)”, and (4) “Latently infected (Resting)” (**Figure 3A**). Of note, the latently infected population is a mixture of uninfected and latently infected cells, which are indistinguishable phenotypically.

**Figure 3.**
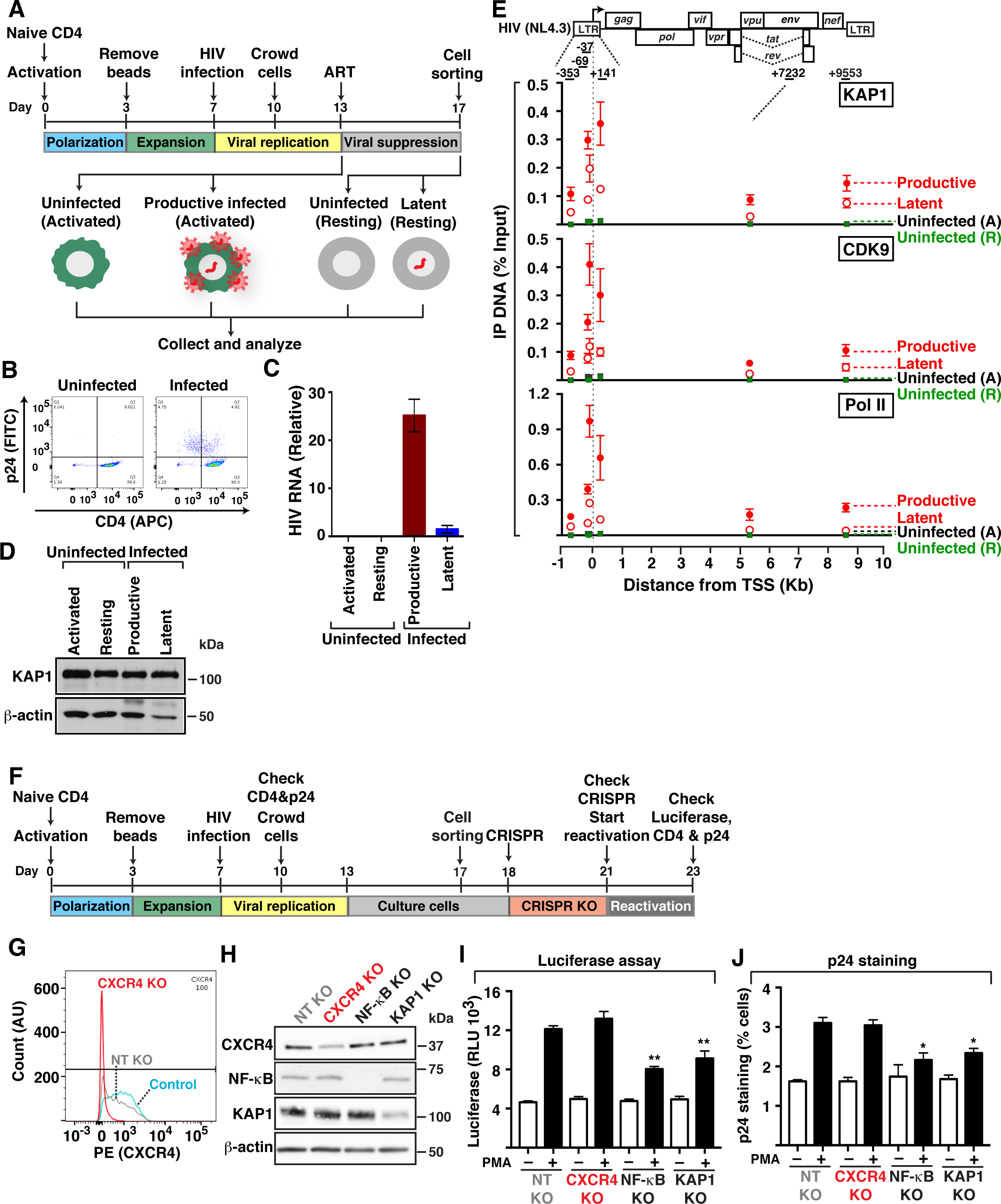
KAP1 Recruitment to the HIV Genome During Productive and Latent Infections in Primary Cells is Required for Provirus Activation. (A) Experimental outline through which naïve cells were used to generate central memory T cells (T_CM_) infected or not with replication competent HIV (HIV-1_NL4.3_) and either activated only or activated and then allowed to transition into a quiescent/resting T cell state. (B) FACS plots (CD4, HIV p24) of uninfected and infected cells as in panel (A). (C) Quantitation of HIV transcripts (using HIV(+7232) amplicon) normalized to *ACTB* by RT-qPCR in the four cell states indicated (mean ± SEM; n =3). (D) Western blots of the four cell states from panel (A) with the indicated antibodies. (E) Top, scheme of the HIV provirus. The arrow denotes the position of the transcription start site (TSS) on the 5’-LTR. Bottom, ChIP assays were performed with protein extracts from the four cells states from panel (A) and the indicated antibodies (KAP1, CDK9, and Pol II) followed by qPCR assays with a series of amplicons mapping throughout the entire provirus (mentioned at the top of the schematic) to monitor factor interactions with the HIV genome. A normal IgG serum was used as a negative control in ChIP-qPCR, which revealed no amplification (not shown). The ChIP-qPCR data was normalized using the “Percent Input Method” (see Experimental Procedures for full description of normalization). Values represent the percentage (%) of Input DNA immunoprecipitated (IP DNA) and are the average of three independent experiments (mean ± SEM; n =3). Note that “Uninfected” refers to both activated (A) and resting (R) T cell state conditions. (F) Experimental outline through which naïve cells were used to generate central memory T cells (T_CM_) that were then infected with replication competent, pseudotyped viruses pNL4.3-delta*Env*-nLuc-2A*Nef*-VSVG and used for CRISPR-Cas9–mediated KO of CXCR4, NF-κB (p65 subunit) and KAP1, to be used in reactivation assays (panels I and J). (G) FACS plots (CXCR4) in control T_CM_ (not nucleofected) and T_CM_ nucleofected with Cas9-gRNA complexes for a non-targeting control (NT) and targeting CXCR4. (H) Western blots of the four CRISPR-Cas9 experiments with the indicated antibodies. (I) Luciferase assay of T_CM_ cells containing KO of specific host factors generated as in panel (F) and treated with PMA or vehicle (DMSO). Luciferase is expressed as relative luciferase units (RLU). (J) p24 staining of T_CM_ cells containing KO of specific host factors generated as in panel (F) and treated with vehicle (DMSO) and PMA (-/+ PMA, respectively). Statistical significance in panels (I) and (J) was determined using unpaired Student’s *t*-test. \**P* < 0.05, \*\**P* < 0.005, \*\*\**P* < 0.0005.

The four cell states were then collected for subsequent analysis of viral gene expression by RT-qPCR, KAP1 expression by western blot, and factor occupancy at the proviral genome by chromatin immunoprecipitation (ChIP) assay (**Figures 3C–E**). Because of the low levels of reactivation in the presence of T cell activation in this model (∼1%), as judged by p24 levels by FACS (Bosque and Planelles, 2009; Martins et al., 2016), we collected productive infection data to circumvent the low levels of latency reversal. FACS analysis confirmed the expectation that uninfected cells displayed high levels of CD4 with no evidence of p24 staining, while infected cells showed a reduction of CD4 levels (there is a broad range of CD4 down-modulation, from wild-type levels to one log; and cells that appear not to down-modulate are early infected, typically, and later transition to overt down-modulation) (**Figure 3B** and data not shown). Consistently, examination of HIV transcript levels indicated that the productively infected state had ∼25-fold higher transcript levels than the latent state; and that, as expected, no transcripts were detected in the uninfected states, whether resting or activated (**Figure 3C**). Given that a low-level of HIV RNA molecules were detected in the latent state, we cannot exclude the possibility that this reservoir is made up of a combination of inactive and low-level transcribed proviruses.

To determine whether KAP1 is expressed in the four cell states we performed western blot and observed that KAP1 is detected at similar levels in both infected and uninfected cells with no, or little, effect of the cell state (active versus resting) (**Figure 3D**). The fact that KAP1 is expressed in both activated and resting primary T_CM_ prompted us to determine whether KAP1 is recruited to the HIV proviral genome during productive and latent infections. Given that in transformed cell-based models (J-Lat) we observed KAP1 recruitment to the proviral LTR (McNamara et al., 2016b), we predicted that latently infected cells would contain KAP1 at the LTR as well. To test this possibility, we performed ChIP assays in the four cell groups and observed that KAP1 along with Pol II and the elongation complex (CDK9) are bound to the LTR both in productively and latently infected cells (**Figure 3E**). In addition, we detected higher levels of KAP1 bound to the proviral 5’-LTR and within the genome in productively infected cells, compared to latently infected cells, probably due to the higher levels of transcription activation during productive infection (as observed with both Pol II and CDK9) (**Figure 3E**), consistent with HIV expression data (**Figure 3C**).

Taken together, we report that: i) KAP1 is expressed in primary T cells irrespective of cell state (active or resting), ii) KAP1 and the host transcription elongation complex is recruited to the proviral genome in the primary T_CM_ model, and iii) the levels of transcription elongation complex recruitment mirror the proviral fate state, with higher levels in the active compared to the latent state.

### CRISPR-Cas9 Reveals a Critical Role for KAP1 in Reactivation of Latent HIV in Primary Cells

Given KAP1 is expressed and recruited to the proviral genome in the primary T_CM_ model, we then wanted to examine whether KAP1 could drive HIV transcription activation of latent proviruses in response to immune cell signaling. To test this, we infected T_CM_ cells with replication incompetent luciferase-tagged virus, and active infections (p24-positive/CD4-negative) were isolated using a magnetic sorting kit to enrich in latent infections (**Figure 3F**) as shown in Figures 3A and B. After expansion of the remaining CD4+ T cells (both uninfected and latently infected), *in vitro* preformed CRISPR-Cas9 ribonucleoprotein (RNP) complexes (Zuris et al., 2015) were delivered into cells to knockout (KO) KAP1. We also included several controls: CXCR4 (a positive cell surface marker that allows for ease of KO visualization by FACS), the NF-κB p65 subunit (a positive control for reactivation assays since it is required for activation of the latent virus in response to immune stimulation), and a negative non-targeting (NT) guide RNA-containing RNP complex not specific for any known human gene.

Remarkably, as revealed by flow cytometry and western blot, the CRISPR/Cas9 RNP-based KO approach was selective and efficient, albeit with different efficiency levels (ranging from ∼50-to-100%) (**Figures 3G and 3H**). After determining KO efficiency, reactivation of latent viruses was then computed using luciferase assays and p24 staining. Notably, we observed that KAP1 KO does not affect levels of proviral activation in the absence of immune cell signaling (-PMA, phorbol ester), similarly to CXCR4 KO and NF-κB KO (**Figures 3I and 3J**), consistent with the idea that KAP1 does not control HIV transcription at steady-state conditions, in agreement with the experiment in **Figure S1I**. However, interestingly, we found that KAP1 KO dampens both luciferase levels and intracellular p24 expression in response to immune cell signaling (+PMA), demonstrating its importance in the host cell response to immune stimulation. Remarkably, this result mirrors the KO of the master regulator NF-κB, which is key for activation of the host phase, because NF-κB KO dampened ∼50% reactivation of latent HIV in response to PMA (**Figures 3I and 3J**). While the levels of reactivation from latency in the primary system are very low (∼3% of total cells) because of the relatively low dynamic range of the assay, it must also be noticed that the reduction in reactivation efficiency after KAP1 KO is ∼50%, a significant effect considering that complete gene KO in primary cells could not be achieved. More importantly, however, this affect approximates in magnitude the effect of NF-κB KO, arguably the most important transcription factor required for latent HIV reactivation, for which more efficient depletion was achieved. Therefore, these data strongly support an important role for KAP1, which is on par with that of the established transcription factor NF-κB.

In contrast with KAP1 and NF-κB KO, CXCR4 KO showed no effects on proviral responses to immune stimulation, despite its efficient depletion levels, thus revealing specificity in the CRISPR-Cas9 results. In addition, importantly, we also observed these results could be recapitulated using the SupT1 CD4+ T cell line (**Figure S2**), both in terms of KO efficiency (∼50-90% depending on the target) as well as decreased reactivation of latent viruses in response to PMA stimulation, thus demonstrating KAP1 (like the master transcriptional regulator NF-κB) has a crucial role in proviral transcription activation and fate in both primary and T cell lines.

### KAP1 is Central for Activation of the Host Phase of the HIV Transcriptional Circuit

The previous data suggested that KAP1 (and the transcription elongation machinery) facilitate activation of the HIV transcriptional circuit in primary cells, but these experiments do not distinguish which phase of the circuit (host or viral) and which step of the transcriptional cycle (initiation or elongation) KAP1 controls. Thus, having established this essential function we then asked: What is the contribution of KAP1 to the different phases of the HIV transcriptional circuit; and how does KAP1 contribute to the positive feedback loop (which is crucial for completion of the viral life cycle and failure to maintain it has been implicated in the establishment of proviral latency)?

As discussed above (**Figure 1**), the HIV transcriptional circuit is composed of two phases: 1) host phase (NF-κB–driven), and 2) viral phase (Tat–driven). Thus, to define in which phase of the HIV transcriptional circuit KAP1 participates, we efficiently silenced (>90%) the expression of KAP1 in two different cell-based models that recapitulate the different phases of HIV transcriptional circuit during infection: (1) The Jurkat HIV Tat+ clone (E4) contains a replication-competent virus that is transcribed by the sequential action of NF-κB and Tat in response to immune stimulation (TNF), and (2) the Jurkat HIV Tat-clone (2B2D) contains a replication-defective virus that is only transcribed by NF-κB in response to TNF because of a non-functional Tat mutant (Tat C22G) (Pearson et al., 2008) (**Figures 4A and 4B**).

**Figure 4.**
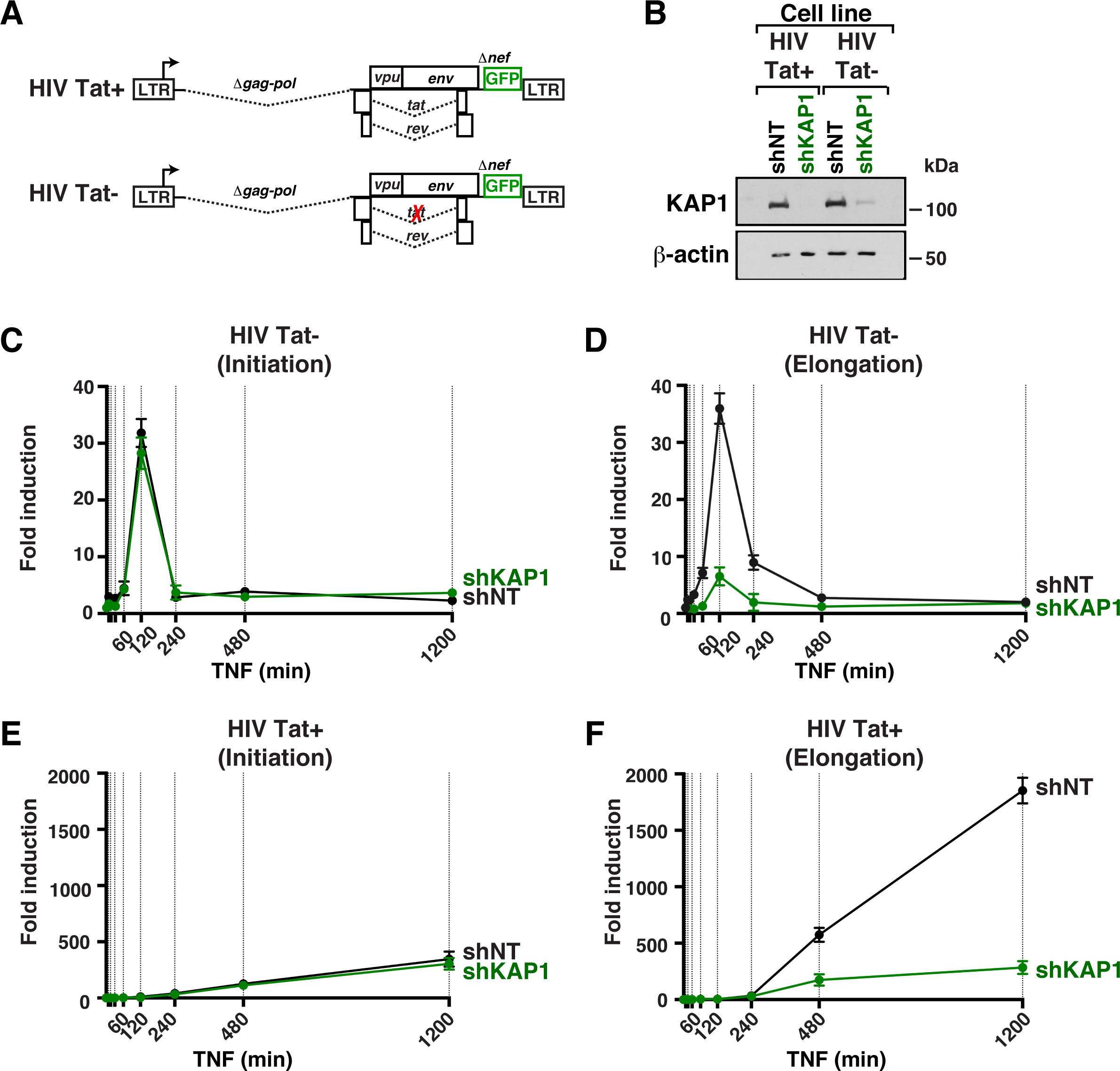
KAP1 is Required for Activation of the Host, but not the Viral, Phase of the HIV Transcriptional Program. (A) Scheme of the HIV Tat+ (E4) and HIV Tat-(2B2D) proviruses containing wild-type Tat or the non-functional Tat C22G mutant (Pearson et al., 2008), respectively. (B) Western blots of the four cell-based models with the indicated antibodies. (C) Fold HIV RNA induction (initiating transcripts) in the HIV Tat-shNT and shKAP1 cell lines in response to a time course TNF treatment. (D) Fold HIV RNA induction (elongating transcripts) in the HIV Tat-shNT and shKAP1 cell lines in response to a time course TNF treatment. (E) Fold HIV RNA induction (initiating transcripts) in the HIV Tat+ shNT and shKAP1 cell lines in response to a time course TNF treatment. (F) Fold HIV RNA induction (elongating transcripts) in the HIV Tat+ shNT and shKAP1 cell lines in response to a time course TNF treatment.

To test the contributions of KAP1 to the host phase (NF-κB–driven) and positive feedback loop (sequential action of NF-κB and Tat) of the HIV transcriptional circuit, we measured temporal HIV expression in the four cell lines in response to TNF, using amplicons that can measure promoter-proximal (indicative of transcription initiation) and promoter-distal (indicative of transcription elongation) transcripts in RT-qPCR assays (**Figures 4C–4F**). We observed that NF-κB similarly activates transcription initiation in the host phase (∼30-fold over untreated cells) in both HIV Tat-cell lines (shNT and shKAP1) (**Figure 4C**). However, loss of KAP1 blunted transcription elongation (∼5.5-fold less elongation in the shKAP1 cell line compared with shNT) (**Figure 4D**), indicating that KAP1 plays an important role in controlling proviral transcription elongation in response to immune stimulation. To analyze the combined effect of KAP1’s loss on both host (NF-κB) and viral (Tat) phases we examined the HIV Tat+ cell lines. Consistent with the previous results, while KAP1 silencing showed a minimal effect on proviral transcription initiation (**Figure 4E**), the largest effect was observed at the elongation step, as revealed by a ∼6.5-fold higher activation levels in the shNT cell line over shKAP1 (**Figure 4F**).

It is worth noting that the magnitude of transcription activation in the two systems is largely different because the HIV Tat-provirus is only activated by NF-κB (∼35-40-fold activation) and the HIV Tat+ provirus is activated by the sequential action of NF-κB and Tat (∼1800-fold activation) (**Figure 1**) leading to full activation of the HIV transcriptional circuit through the feedback loop (compare plots in **Figures 4D and 4F**). This is why host phase activation follows a unimodal distribution and Tat activation shows an exponential pattern of activation (at least in the times examined), consistent with an auto-activation model as in the positive feedback loop (Razooky and Weinberger, 2011; Weinberger et al., 2005).

Taken together, the data indicate that the host phase strictly relies on KAP1 function. Given the fragility of the host phase (due to loss of KAP1), the magnitude of the Tat feedback gets compromised.

### Tat Functions in a KAP1–independent Manner to Facilitate Transcription and Reactivation of a Latent Virus by Directly Recruiting the Host Elongation Complex to the HIV Genome

The previous data suggested a model in which KAP1 functions as a transcriptional co-activator of NF-κB to activate the host phase of the HIV transcriptional circuit in response to immune cell signaling. Given KAP1 plays a critical role in transcription activation of the host phase thereby influencing the magnitude of the feedback loop, none of the cell-based systems previously used allowed us to directly test the role of KAP1 on activation of the viral phase. Thus, to directly examine the functional role of KAP1 in this context, we co-transfected U2OS shNT and shKAP1 cell lines with an LTR firefly (FFL) LUC reporter, increasing amounts of Tat, and a constitutive CMV-Renilla (RL) as internal control, and calculated the FFL/RL ratio as previously described (D’Orso et al., 2012). We observed that Tat similarly activates both cell lines in a dose-dependent manner (**Figure 5A**), irrespective of the high efficiency of KAP1 KD (**Figure 5B**), indicating that Tat can function, at least in this reporter assay, in a KAP1-independent manner.

**Figure 5.**
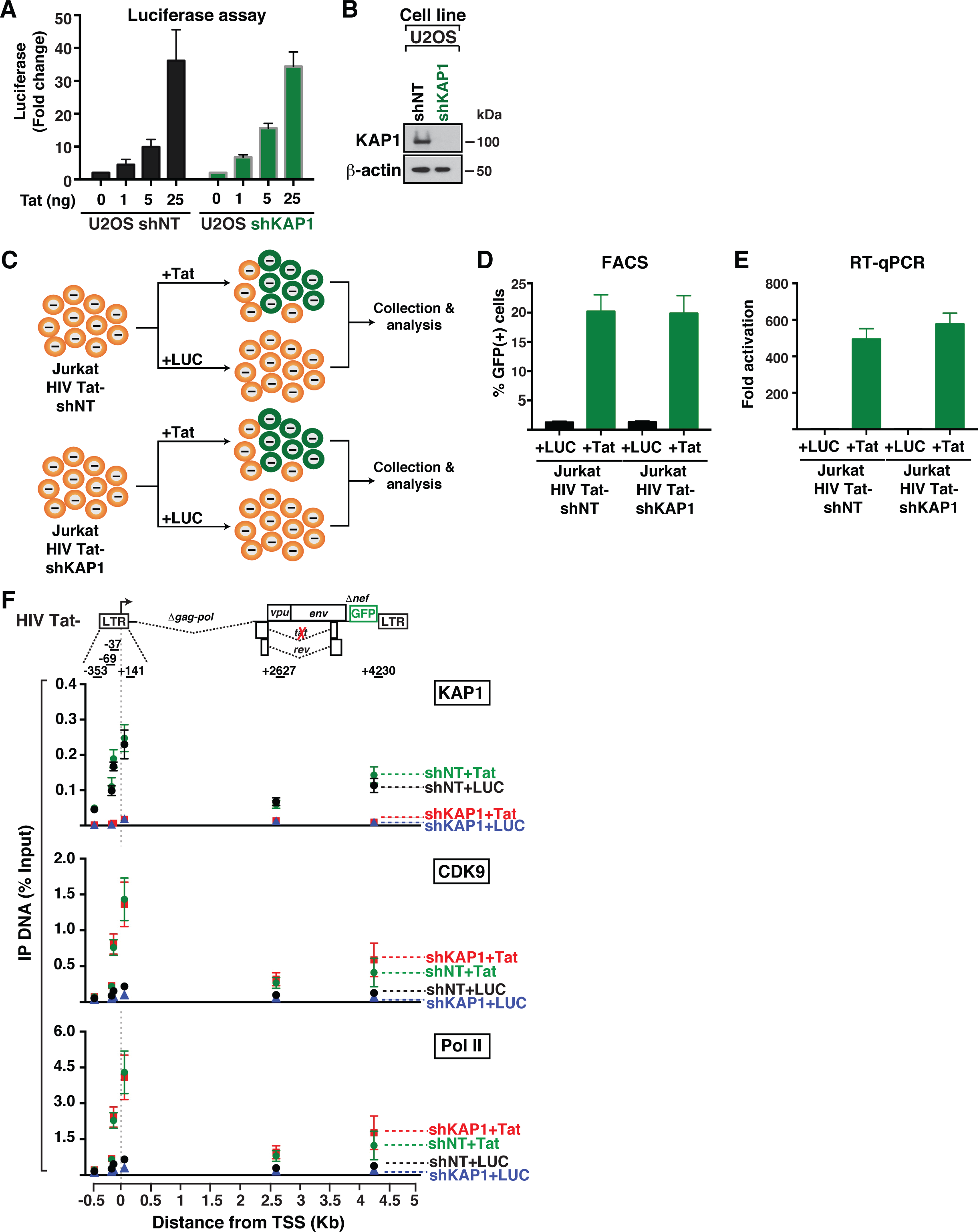
The Master Regulator of the Viral Phase (Tat) Operates in a KAP1-independent Manner. (A) Quantification of firefly luciferase activity (FFL) from an HIV LTR-FFL reporter in the absence (0) or presence of increasing Tat concentrations (normalized to a constitutive CMV-Renilla (RL)) in the shNT and shKAP1 U2OS cell lines. (B) Western blots of the indicated U2OS cell lines used in panel (A). (C) Experimental outline in which the HIV Tat-shNT and shKAP1 cell lines were transduced with pTRIP-LUC or pTRIP-Tat lentiviruses at day 1 and collected at day 3 for their analysis by FACS and RT-qPCR assays (panels (D) and (E), respectively). (D) Flow cytometry based quantitation of the percentage of GFP+ cells after transduction of the shNT and shKAP1 cell lines from panel (C) with the indicated lentiviruses expressing LUC and Tat. (E) RT-qPCR assay of RNA isolated from the cell lines from panel (C) using the elongation amplicon (+2627). The change in HIV gene expression is shown as fold HIV RNA activation (Tat over LUC). (F) Top, scheme of the HIV Tat-(2B2D) provirus. The arrow denotes the position of the transcription start site (TSS) on the 5’-LTR. Bottom, ChIP assays were performed with protein extracts from the four cells states from panel (C) and the indicated antibodies (KAP1, CDK9, and Pol II) followed by qPCR with a series of amplicons mapping throughout the entire provirus (indicated at the top of the schematic) to monitor factor interactions with the HIV genome. A normal IgG serum was used as a negative control in ChIP-qPCR, which revealed no amplification (not shown). The ChIP-qPCR data was normalized using the “Percent Input Method” (see Experimental Procedures for full description of normalization method). Values represent the percentage (%) of Input DNA immunoprecipitated (IP DNA) and are the average of three independent experiments (mean ± SEM; n =3).

If Tat efficiently transactivates the provirus bypassing KAP1 then the result should be independent of the cell-based model used. To test this idea, we transduced the cell lines containing Tat-defective proviruses (Jurkat HIV Tat-shNT and HIV Tat-shKAP1) with lentiviruses expressing Tat and LUC (as negative control) (**Figure 5C**). First, to make sure transduction was efficient, we collected cells at day 3 post-transduction to quantitate the number of GFP+ cells by FACS (**Figure 5D**) and to measure HIV gene expression by RT-qPCR (**Figure 5E**). Notably, we observed that Tat transduction increases the levels of GFP+ cells by ∼20-fold over transduction with LUC-expressing lentiviruses (**Figure 5D**), in agreement with the robust synthesis of viral transcripts in the presence of Tat (∼550-fold over LUC) (**Figure 5E**). Notably, transduction of the non-functional Tat mutant C22G is not able to restore transcription activation (**Figure S3**). Together this data supports the notion that ectopic Tat can function in a KAP1-independent manner to trans-activate the integrated provirus (viral phase), in agreement with the reporter assay (**Figure 5A**).

If the viral phase functions in a KAP1-independent manner, we would expect Tat to recruit CDK9 and promote Pol II function in the absence of KAP1. To test this model, we performed ChIP assays to measure the occupancy of KAP1, CDK9 and Pol II throughout the HIV genome (promoter and gene body) in the Tat-defective Jurkat cells (HIV Tat-) in four different scenarios: with/without KAP1 and with/without Tat (**Figure 5F**). By comparing the occupancy levels of CDK9 and Pol II in the absence/presence of KAP1 with Tat (shKAP1+Tat and shNT+Tat, respectively) we can interrogate whether Tat function requires KAP1. In addition, by comparing the levels of CDK9 and Pol II in the absence/presence of KAP1 without Tat (shKAP1+LUC and shNT+LUC, respectively) we can examine changes occurring at the basal level (without immune cell signaling).

First, ChIP assays revealed that loss of KAP1 in the absence of Tat reduced CDK9 recruitment (∼2.2-fold) to the promoter, consistent with previous data (McNamara et al., 2016b). Second, interestingly, Tat recruits CDK9 to the HIV promoter and inside the proviral genome with similar efficiencies, irrespective of the presence of KAP1 (**Figure 5F**). Consequently, Tat promotes the recruitment of more Pol II to the viral promoter and significantly enhances the levels of Pol II inside the proviral genome, consistently with its role in promoting transcription elongation. The data suggest that Tat (viral phase) functions in a KAP1-independent manner to recruit the elongation complex to the provirus to promote its activation.

Collectively, the data of Figures 4 and 5 demonstrate that the two phases of the HIV circuit (host and viral) have different functional requirements, and that HIV may have evolved Tat to circumvent the KAP1-regulated step to directly recruit the kinase to the proviral genome and sustain transcription. We propose that this ‘minimalist’ regulatory system that HIV evolved might explain why Tat functions as a potent transcription activator compared with cellular activators, and why its function is critical for efficient viral replication as well as reactivation of latent viruses. Despite these discoveries, we are not completely ruling out the possibility that Tat could cooperate with KAP1 in other scenarios. Given HIV can integrate semi-randomly, it is possible that in different integration sites, depending on the chromatin environment, Tat may need KAP1 for proviral activation through currently unknown mechanisms. This opens an interesting scenario for future investigations.

### Mathematical Modeling the HIV Transcriptional Circuit Reveals a Critical Function for KAP1 in Modulating the Feedback Loop Thus Shaping Proviral Fate

HIV is efficiently activated in response to immune cell signaling through the sequential action of host (NF-κB) and viral (Tat) activators (Karn, 2011; Nabel and Baltimore, 1987). In summary, our data revealed two critical phases of the HIV transcriptional circuit. First, in the host phase, NF-κB rapidly translocates from its latent, cytoplasmic state into the nucleus where it binds the proviral promoter, eliciting KAP1-dependent CDK9 activity, productive transcription elongation, and Tat synthesis. Second, in the viral phase, Tat promotes the KAP1-independent feedback loop to robustly activate HIV transcription thereby promoting viral replication (**Figure 6A**).

**Figure 6.**
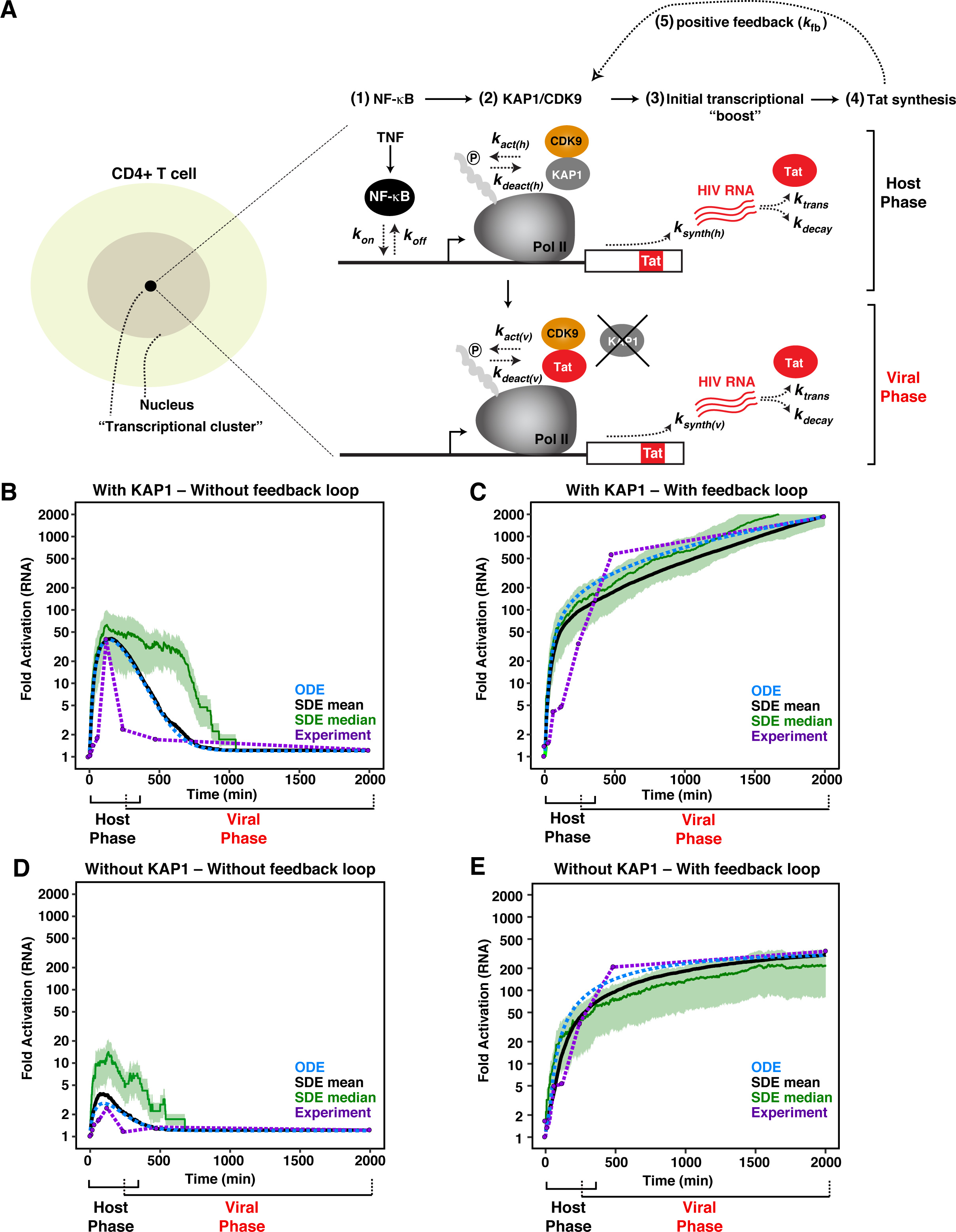
Mathematical Model of a Complete HIV Transcriptional Program. (A) Left, scheme of a CD4+ T cell containing a transcriptional cluster. Right, simplified model of the host (NF-κB–driven) and viral (Tat–driven) phases of the HIV transcriptional program. α is the rate of basal HIV RNA synthesis (not shown for simplicity). The model parameters are shown as: (1) *k*_*on*_, *k*_*off*_, the rate of NF-κB association and dissociation from the promoter in response to TNF stimulation; (2) *k*_*act(h*)_, *k*_*deact(h)*_, the rates of KAP1-mediated recruitment and activation/deactivation of CDK9 in the host phase; (3) *k*_*synth(h)*_, the rate of HIV RNA synthesis by NF-κB (initial transcriptional boost; host phase); *k*_*trans*_, *k*_*decay*_, the rate of Tat RNA translation and decay in the absence of feedback loop; (4) *k*_*act(v)*_, *k*_*deact(v)*_, the corresponding rates of Tat–CDK9 association and dissociation from the promoter; *k*_*synth(v)*_, the rate of HIV RNA synthesis by Tat (feedback loop; viral phase); (5) *k*_*fb*_, the rate of Tat positive feedback loop induced by Tat recruiting new CDK9 to the promoter and activating transcription elongation. (B) Computational simulation of HIV RNA synthesis (fold activation) in cells normally expressing KAP1 and infected with HIV Tat-(without feedback loop) in the presence of immune stimulation (TNF) over untreated cells. (C) Computational simulation of HIV RNA synthesis (fold activation) in cells normally expressing KAP1 and infected with HIV Tat+ (with feedback loop) in the presence of immune stimulation (TNF) over untreated cells. (D) Computational simulation of HIV RNA synthesis (fold activation) in cells lacking KAP1 expression and infected with HIV Tat-(without feedback loop) in the presence of immune stimulation (TNF) over untreated cells. (E) Computational simulation of HIV RNA synthesis (fold activation) in cells lacking KAP1 expression and infected with HIV Tat+ (with feedback loop) in the presence of immune stimulation (TNF) over untreated cells. For panels B–E, the y-axis was set to log_10_ scale to allow better comparisons among all four simulations. Black curve denotes stochastic differential equations (SDE) mean. Blue dotted curve denotes ordinary differential equations (ODE). Green curve denotes the SDE median. Purple dotted line and purple data points denote experimental data. Note that the duration of two phases of the circuit is an approximation based on the length of the host phase, when the host phase starts decaying in HIV Tat-proviruses, and when the viral phase starts progressing before the decay of the host phase in cells infected with HIV Tat + proviruses.

To determine whether these previous findings can be integrated into a theoretical framework of host-viral transcriptional regulation, we developed a mathematical model that describes the minimal set of interactions in a transcriptional system (see **Extended Experimental Procedures** for a detailed description of the model assumptions, parameters, and variables). Previous stochastic computational modeling has revolved around the idea that the HIV transcriptional circuit is composed of two phases: 1) basal (Tat-independent phase), and 2) viral (Tat-dependent) (see **Figure 1E**) (Weinberger et al., 2005). However, as explained above, the complete HIV circuit is composed of three phases: 1) basal, 2) host, and 3) viral (**Figure 1F**). Thus, the contributions of the host phase (which is critical because it primes HIV for activation, as we have experimentally observed (**Figure 4**)) to the feedback loop was not integrated into previous studies. We thus developed a mathematical model that enables one to investigate the individual contributions of the host and viral phases to the HIV transcriptional circuit and thus have a complete view of the real system.

In theory, our model is based on the principle that spontaneous proviral transcription activation in response to immune simulation results from the stochastic fluctuations of key host cell factors (such as KAP1) between the nucleoplasm and promoter interactions in chromatin territories. We propose that the probability of proviral activation is dependent on the coincidence of two events that might occur independently or simultaneously. First, NF-κB must associate with the viral promoter [**Figure 6A**, point (1)]; and second, a KAP1 molecule must bind near the Pol II complex/promoter [**Figure 6A**, point (2)]. Once recruited, KAP1 delivers primed CDK9 for ‘on site’ activation (McNamara et al., 2016b). The active kinase then phosphorylates its substrates (e.g., Pol II) at the promoter [**Figure 6A**, point (2)]. Collectively, this sequence of molecular events initiates proviral transcription and Tat synthesis [**Figure 6A**, point (3)], further recruiting more kinase but, in this case, bypassing the KAP1-centric host cell regulatory system [**Figure 6A**, point (4)], thereby promoting the feedback loop by increasing the number of elongating Pol II molecules [**Figure 6A**, point (5)].

In our theoretical analysis, we considered four conditions with two main variables to model HIV RNA synthesis in response to immune cell signaling:

1. Cells with normal KAP1 expression and feedback loop;
2. Cells with normal KAP1 expression but lacking feedback loop;
3. Cells with loss of KAP1 expression but normal feedback loop; and
4. Cells with loss of KAP1 expression and lacking feedback loop.

Literature values were used to estimate the rates of basal HIV RNA synthesis (**α)**, formation and dissociation of the NF-κB–DNA complex in the host phase (*k*_*on*_, *k*_*off*_), KAP1-mediated recruitment and activation/deactivation of CDK9 in the host phase (*k*_*act(h)*_, *k*_*deact(h)*_), NF-κB–mediated HIV RNA synthesis in the host phase (*k*_*synth(h)*_), Tat-mediated recruitment and activation/deactivation of CDK9 in the viral phase (*k*_*act(v)*_, *k*_*deact(v)*_), Tat-mediated HIV RNA synthesis in the viral phase (*k*_*synth(v)*_), RNA translation (*k*_*trans*_), RNA decay (*k*_*decay*_), and Tat positive feedback (*k*_*fb*_). Furthermore, experimental data was used to calibrate unknown kinetic rates (see **Extended Experimental Procedures**).

Computational simulations resulted in an initial “boost” of TNF-induced NF-κB–mediated HIV RNA synthesis (host phase) in the presence of KAP1, but lack of feedback loop due to Tat’s absence in the system (**Figure 6B**). NF-κB activated ∼50-fold *in silico*, a value that closely resembles the measured NF-κB activation rates (**Figure 4**), even though the magnitude of activation is directly proportional to TNF concentration (see below), consistent with previous data (Tay et al., 2010). In addition, notably, the model gave rise to temporally decay of NF-κB activation in the absence of feedback loop, as is observed *in vivo* (**Figure 4**), and HIV RNA levels return to the low steady-state level of basal transcription (**Figure 6B**). In the presence of normal KAP1 levels and feedback loop (Tat+ provirus), the initial boost is largely amplified by Tat activity leading to an exponential increase (>100-fold activity) (**Figure 6C**), consistent with experimental data (**Figure 4**) and previous studies (Feinberg et al., 1991; Laspia et al., 1989).

In cells lacking normal KAP1 levels, NF-κB activity (host phase) is largely compromised (see the virtual decrease in the initial transcriptional boost) (**Figure 6D**), consistent with the experimental data (**Figure 4**). With the loss of HIV RNA synthesis by NF-κB in cells lacking KAP1, some level of expression can still be observed that is accelerated by the feedback loop, albeit at a much lower rate compared to the KAP1-positive scenario (fold differences of ∼2.8 at *t* = 480 min and ∼5.5 at *t* = 2000 min during the exponential growth phase) (**Figure 6E**).

Together, these data indicate that the stochastic assembly of transcription elongation complexes at the proviral promoter is required to establish the initial proviral transcriptional “boost”. Consistent with this interpretation, NF-κB was unable to activate in the absence of KAP1, despite its efficient binding to the promoter (McNamara et al., 2016b). Remarkably, our mathematical model recapitulates the normal transcription activation pattern of a complete HIV circuit.

### Perturbation analysis of the model and model behavior

We then investigated the behavior for the model during perturbation of model parameters. For this purpose we used the well-mixed deterministic Ordinary Differential Equation (ODE) model, as we were interested in the overall (mean) behavior, disregarding any noise and stochastic fluctuations. As expected in the case of chemical mass action systems (Hahl and Kremling, 2016), both the ODE and the Stochastic Differential Equation (SDE) mean show similar behavior (**Figures 6B–E**). It has been shown that, in linear systems, the mean after SDE and the deterministic variable of the ODE coincide (Hahl and Kremling, 2016). However, skewed fluctuations through large bursts may lead to a shift of stochastic modes away from the mean. Such bursts have not been observed in our stochastic simulations to the extent of causing deviation between the ODE model and the SDE mean. Thus, for simplicity, we utilized the corresponding ODE model to investigate the effect of KAP1 levels on the dynamic behavior. Notably, **Figures 7A** and **7B** capture the experimental measurements and ODE model behavior in the four conditions (with/without KAP1 and with/without feedback loop; see corresponding **Figures 6B-E** with additional trajectories from SDE simulations), thus demonstrating good agreement between experimental data and simulation (ODE and SDE mean) calculations.

**Figure 7.**
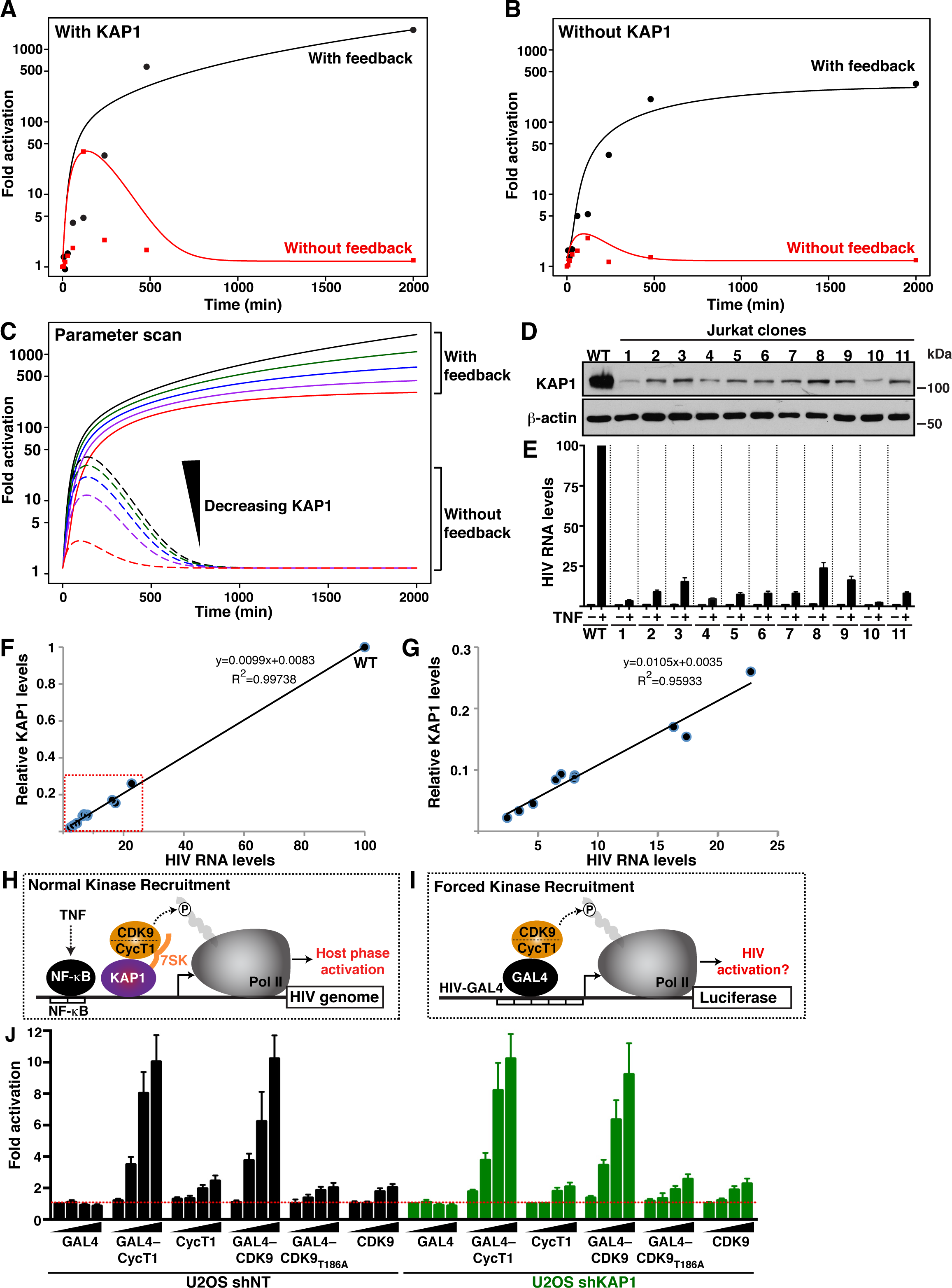
KAP1 Protein Levels Influence the Outcome of the Host Phase Thereby Affecting the Viral Feedback Loop Potential. (A) Computational simulations (ODE) of HIV RNA synthesis (fold activation) in cells normally expressing KAP1 and infected with HIV Tat+ (With feedback, black line) or HIV Tat-(Without feedback, red line) in response to a temporal immune stimulation (TNF) treatment. (B) Computational simulations (ODE) of HIV RNA synthesis (fold activation) in cells lacking KAP1 and infected with HIV Tat+ (With feedback, black line) or HIV Tat-(Without feedback, red line) in response to a temporal immune stimulation (TNF) treatment. (C) Parameter scan computational simulations (ODE) with variable KAP1 levels. The black and red lines correspond to the highest and lowest KAP1 protein levels, respectively. The outcome of the host phase (Without feedback) and host-viral phases (With feedback) is indicated. (D) Western blots of parental (WT) Jurkat HIV Tat+ (E4), and KAP1 knockdown (KD) single-cell sorted clones expressing variable levels of KAP1 protein. (E) Relative HIV RNA expression levels of HIV Tat+ (E4) KAP1 KD clones in the absence (-) and presence (+) of TNF (25 ng/ml) stimulation for 16 hr measured by RT-qPCR (elongation amplicon, +2627). The data of the parental HIV Tat+ (E4) in the absence of TNF was set to 1 and the stimulation to 100 to allow for easier comparisons. Comparison of KAP1 protein levels among the different clones (panel D) and HIV RNA levels in response to immune stimulation (panel E) shows a direct correlation between KAP1 protein levels and HIV proviral transcription. (F) Correlation plot between relative KAP1 protein levels (normalized to levels in E4 = 1) and HIV RNA expression levels in the presence of TNF stimulation. A trend line was fitted to the data points and the Pearson’s correlation coefficient (R^2^) was calculated. (G) Correlation plot inset (red box from panel F) showing all data points except WT cells for easier visualization of datasets in the Jurkat clones examined in panels (D) and (E). (H) Simplified scheme for the KAP1-mediated recruitment of the 7SK-bound P-TEFb kinase (CycT-CDK9) to the HIV promoter to induce Pol II phosphorylation and host phase transcription activation in response to immune stimulation (TNF) by NF-κB. (I) Scheme of a minimal proviral promoter transcribing a Luciferase reporter gene (HIV-GAL4) in which the P-TEFb kinase (CycT-CDK9) is artificially recruited through a heterologous interaction (yeast GAL4) to a minimal HIV promoter to test whether kinase recruitment is necessary and sufficient for gene activation in the absence of the normal recruitment mechanism as shown in panel (A). (J) Fold activation luciferase levels of the minimal HIV-GAL4 reporter from panel (I) transfected with the indicated activators in the U2OS shNT and shKAP1 cell lines from **Figure 5B**.

We then performed a parameter scan by simulating the model multiple times each time varying the value of one parameter, in this case KAP1 protein levels. Strikingly, the parameter scan model revealed a direct relationship between the rate of proviral initial “boost” of transcriptional activation and the levels of KAP1 protein expressed, and therefore potential binding to the proviral promoter (τ[KAP1]) (**Figure 7C**). Although the phenotype is first observed on the initial host-dependent “boost,” it ultimately impacts on the strength of the feedback loop. Here, four states of decreasing host phase activation have proportional effects on the feedback loop. The lower the magnitude of the host phase, due to reduced KAP1 levels, the lower the magnitude of the feedback loop (**Figure 7C**). Together, this signifies that the threshold of activation of the host phase generates stochasticity potentially causing large cell-to-cell differences in transcript levels in patient samples thereby affecting latency reversal potential (see below).

### Testing the Influence of Host Phase Heterogeneity and Immune Cell Signaling Strength on Transcriptional Fragility and Viral Phase Outcomes

The previous simulations indicated that oscillations of KAP1 protein levels during infection could generate cell-to-cell differences thereby creating transcriptional noise in the host phase and impacting homogeneous responses to immune cell signaling and latency-reversing agents (LRAs) due to alterations of the feedback loop (**Figure 7C**). To test this model prediction experimentally, HIV RNA synthesis was monitored over time in response to TNF on several Jurkat HIV Tat+ clones (created through KAP1 KD and single-cell sorting), which express variable KAP1 protein levels as revealed by western blot analysis (**Figure 7D**). We observed that the lower KAP1 protein levels the larger the reduction in HIV RNA levels in response to TNF stimulation (**Figure 7E**). Interestingly, correlation analysis provides direct evidence that HIV RNA levels produced in response to immune cell signaling (TNF) are directly proportional to KAP1 protein levels in the system (**Figures 7F and G**). These results are consistent with the theory that KAP1 amplifies the initial transcriptional “boost” and sustains the host phase (NF-κB activation in response to immune stimulation) thereby impacting on the outcome of the viral phase.

Given that the previous assay used different clonal cell lines, we then created an isopropyl-β-D-thio-galactoside (IPTG)-inducible KAP1 knockdown Jurkat cell line (HIV Tat-shKAP1), as well as a shNT negative control, to better model the dosage effect of KAP1 protein levels on host phase activation in response to immune cell signaling stimulation in the same system (**Figure S4A**). Remarkably, we observed that a dose-dependent reduction of KAP1 protein levels proportionally reduced proviral activation in response to TNF without affecting basal levels (**Figure S4B**), consistent with the idea that KAP1 is required for immune cell signaling, but not basal, transcription. In addition, a time course of TNF-mediated activation of the host phase in the inducible KAP1 KD system showed a ∼6-fold reduction in the magnitude of proviral activation (**Figure S4C**), consistent with the data in the non-inducible system (**Figure 4**). Together, the data reinforce the idea that KAP1 is a key regulator of the host phase and that KAP1 protein levels correlate with the transcriptional magnitude of the host phase and feedback loop.

The HIV transcriptional program appears to function as an ‘off-on’ switch (Aull et al., 2017; Karn, 2011) where in the absence of environmental stimulation the system remains in the ‘off’ state and upon activation is turned ‘on’. However, immune stimulation strength can affect the threshold of activation (Tay et al., 2010) and thresholds allow biological systems to respond based on inputs strengths and are often a key design feature of gene-regulatory circuits (Aull et al., 2017; Karn, 2011). Given this knowledge, we hypothesized a model in which variable levels of TNF stimulation should generate ‘on’ states with different thresholds, with a concomitant reduction in host/viral phase activation levels in the absence of KAP1.

To test this model, we compared the levels of HIV RNA synthesis produced by HIV Tat-proviruses in response to three TNF inputs (high, medium, and low) and observed the graded decrease of HIV RNA signal intensity in control cells (shNT) as a function of reduced immune stimulation strength (**Figure S4D**). Furthermore, we detected proportionally reduced RNA synthesis in the shKAP1 cell line compared to shNT (**Figure S4D**), indicating that both KAP1 protein levels and stimulation input strength control HIV provirus transcriptional output in the host phase. Similarly, we observed that the reduced HIV RNA synthesis in response to a decreased graded TNF levels in the host phase, directly impacts the magnitude of feedback loop activation in HIV Tat+ proviruses, and again much reduced levels (∼4-6–fold) in the shKAP1 compared to shNT cell lines (**Figure S4E**).

Given that TNF is a strong immune stimulus, we then asked whether known LRAs that function through different mechanisms such as Bryostatin (PKC agonist) and suberoylanilide hydroxamic acid-SAHA-(pan-histone deacetylase inhibitor) would show similar KAP1-dependent activation mechanisms. We thus treated the Jurkat HIV Tat- and HIV Tat+ shNT and shKAP1 cell lines with Bryostatin o SAHA and observed that loss of KAP1 also affected latency-reversal of the host phase and impacts on the threshold of activation of the viral phase, respectively (**Figures S4F and G**), implying that KAP1 plays a key role in HIV proviral activation of the host phase in response to strong immune modulators as well as commonly used LRAs.

Collectively, the data suggest the host phase is subject to tight control by host cell co-factors (like KAP1) whose activity is indispensable to maintain active proviral transcription and avoid the establishment of latency. Conversely, HIV evolved a minimalist system whereby Tat functions in a KAP1-independent manner, bypassing the requirement of the host cell checkpoints to sustain the viral elongation program. Importantly, our experimental and mathematical modeling indicates that fluctuations in KAP1 affect the initial transcriptional burst thereby disturbing the feedback loop and the latency reversal potential.

### Forced Elongation Complex Recruitment to the HIV Promoter Bypasses KAP1 Requirement for Transcription Activation, Like in the Viral Program

Our previous studies suggested KAP1 enables recruitment of 7SK RNA-bound CDK9 to the proviral genome to promote NF-κB–dependent transcription elongation in response to immune stimulation (“Normal Kinase Recruitment”) (**Figure 7H**). Loss of KAP1 could then abolish transcription elongation because CDK9 is not properly recruited to the HIV promoter despite normal NF-κB binding kinetics (McNamara et al., 2016b). If KAP1 functions as a transcriptional co-activator of NF-κB by recruiting CDK9 to the promoter in response to immune stimulation, then we would expect that forced CDK9 recruitment to the promoter (“Forced Kinase Recruitment”) (**Figure 7I**) should facilitate transcription activation in a KAP1-independent manner, bypassing KAP1’s role as co-activator, and functioning in a manner similar to Tat (like in the viral program).

We thus sought to test this predictive model by delivering P-TEFb directly to the promoter through the heterologous yeast GAL4 DNA-binder (**Figure 7I**) (Southgate and Green, 1991). In this context, GAL4 mimics KAP1 to deliver the kinase to the promoter for transcription activation. To test this model, we used a U2OS cell line where KAP1 has been efficiently KD (shKAP1) (**Figure 5B**), or a non-targeting shRNA control cell line (shNT), to transfect a minimal promoter containing 5 copies of the GAL4 binding site fused to FFL LUC (GAL4-LUC reporter) (Southgate and Green, 1991), along with increasing amounts of GAL4, and GAL4-fused or unfused P-TEFb subunits (CycT1 and CDK9) plasmid DNAs. Notably, we found that GAL4–CycT1 similarly activates the HIV reporter in both the NT and KAP1 KD cell lines, with no or minimal effect of GAL4 or unfused CycT1 (**Figure 7J**). Supporting this data, fusion of GAL4 to the catalytic subunit of P-TEFb (CDK9), but not a non-functional kinase (CDK9 T186A mutant) (Li et al., 2005) or the unfused CDK9, largely activates the viral promoter despite KAP1 expression (**Figure 7J**), indicating that, at least in this artificial system, forced recruitment of CDK9 to a promoter bypasses the requirement for KAP1 toward transcription activation.

Collectively, the data indicate that one key function of KAP1 is to recruit CDK9 to the promoter for viral activation. By rewiring the circuit to operate through promoter-bound CDK9, KAP1 becomes dispensable for activation, like in the viral phase of the program. However, since the reporter system used is artificial, this result does not provide quantitative evidence that KAP1 scales with CDK9 recruitment. Additionally, the result does not rule out KAP1 could play other essential roles in the transcriptional cycle, which open interesting scenarios for future research.

## DISCUSSION

Viruses have evolved unique strategies to regulate gene expression, rewire host cell programs, and evade immune system responses to survive within their hosts (Beachboard and Horner, 2016; Reeder et al., 2015). These strategies include a broad range of functions including the use of internal ribosome entry sites (IRES) to hijack host cell machinery for viral RNA translation, use of unique proteins that bind the cap RNA to protect their RNA molecules and regulate their translation, as well as use of a diverse set of restriction factors to evade host antiviral responses (Beachboard and Horner, 2016; Dickson and Wilusz, 2011; Yamamoto et al., 2017). In this work, we found that HIV hijacks the host transcription elongation complex to regulate transcription of its genome in a unique way. Specifically, we discovered that HIV evolved a minimalist circuit bypassing host regulatory checkpoints to robustly active transcription, thus providing another example on how viruses fuel molecular innovation for their own benefit.

For over two decades, it was assumed the host and virus utilize similar regulatory strategies to stimulate transcription from the proviral genome. In these previous models, cellular and viral activators were proposed to recruit CDK9 (as part of the transcription elongation complex) to the proviral genome to facilitate Pol II pause release and transcription elongation (Peterlin and Price, 2006). Here, we demonstrate that while the two phases require CDK9 for activation, the kinase is recruited differently. In the host, the KAP1 co-activator assists the process of transcription by cellular activators by directly recruiting CDK9 to the promoter. In contrast, HIV evolved Tat, which represents a switch to a ‘higher gear’ bypassing KAP1. Interestingly, despite the minimalistic nature of the viral phase, malfunction of the host phase (which primes HIV for activation) directly impacts on the extent of the Tat feedback loop and the transcriptional program as a whole suffers from host “fragility”. We call this phenomenon “transcriptional circuit fragility”. This new notion proposes that activation of the transcriptional circuit in the host phase does not follow a “deterministic” trajectory and experiences stochastic outcomes. This concept differs from the previously proposed stochastic variability (gene expression “noise”) described for the circuit in the basal state (Ho et al., 2013; Rouzine et al., 2014; Weinberger et al., 2005).

When a cell in the basal state is exposed to an environmental cue from the immune microenvironment (e.g., a cytokine), such cue turns on a signaling cascade leading to activation of several molecular events including translocation/activation of latent transcription factors from the cytoplasm to the nucleus, degradation of inhibitory components of the cascade, recruitment of transcriptional co-activators (e.g. KAP1 and CDK9) to chromatin, post-translational modifications on histone tails and other non-histone proteins, increased retention time of transcriptional activators on chromatin, and increased Pol II recruitment to promoters, as well as release from the promoter-paused state. As such, the concept of transcriptional circuit fragility implies that the regulatory program of the host phase is ‘fragile’ meaning that protein level fluctuations or malfunction of key factors implicated in one or more of the molecular events above mentioned would generate an abnormal activation threshold in the program leading to variable cell-to-cell transcriptional outcomes. Our work provides a prime example with KAP1 as key regulator of the elongation switch to facilitate normal activation of the HIV transcriptional program.

To study the concept of transcriptional fragility in more detail and provide a unique resource for the field, we created a mathematical model that accurately predicts the contribution of the host and viral phases of the HIV transcriptional program in response to immune stimulation. Notably, this model recapitulates the experimental data and reveals a critical function for KAP1 (and the transcription elongation complex) in controlling the host, but not the viral, phase of the HIV circuit. As such, loss of KAP1 blunts the initial transcriptional boost thereby dampening the viral phase (feedback loop) and the magnitude of reactivation of a latent virus, thus contributing to the proposed “fragility” model.

Using the *ex vivo* primary model of latency in T_CM_, we demonstrate that: i) KAP1 is expressed in both resting and activated T cell states, ii) KAP1 alongside the elongation complex (CDK9) bind the proviral promoter in both states, and iii) KAP1 contributes to proviral reactivation in response to immune cell signaling, thus providing critical insights for the function of this transcriptional program in a physiologically relevant system.

Previous studies have reported P-TEFb levels (CycT1 and CDK9 T-loop phosphorylated) are very low in resting, non-dividing CD4+ T cells (naïve and memory) in comparison with activated, dividing cells, thus potentially compromising 7SK snRNP complex formation (Budhiraja et al., 2013). However, the low levels of P-TEFb in the resting state do not really limit its sequestration into the 7SK complex, since we have shown that the transcription elongation complex (KAP1 and CDK9) can be recruited to the proviral promoter in the resting state to allow for a much lower level of proviral transcription. It makes complete sense activated cells would have higher levels of competent P-TEFb/7SK than resting cells because they have to cope with a much higher transcriptional demand (activated cells transcribe a more diverse set of coding/non-coding regions than resting cells (Michel et al., 2017; Teague et al., 1999)). This idea is consistent with previous data suggesting the 7SK complex is not just a passive inhibitory entity but contributes to deliver primed kinase to target gene promoters to time the transition from Pol II pausing to pause release (McNamara et al., 2016b). Thus, levels of the competent KAP1-containing elongation complex in the primary system appear to be proportional to the level of cell activation state and its transcriptional requirements.

Even though our data clearly favors a model in which KAP1 is critical for activation of the host phase, we are not formally excluding the involvement of additional co-factors including master regulators, co-activators, and pioneer factors required for chromatin accessibility, among others. So by no means we are proposing that KAP1 is the “major factor” but a key co-activator of the host phase because it helps relieve the elongation blockage at the promoter through recruitment of primed CDK9. As such, importantly, the concept of transcriptional fragility is generalizable, implying that malfunction/dysregulation of any other key component required for activation of the host phase, including the most relevant master regulators that recognize *cis*-elements at the viral promoter (i.e., Sp1, NF-κB, NFAT, and AP-1), co-activators such as histone acetyl transferase, kinases, and phosphatases, and/or promote chromatin remodeling (i.e., BRD4, BAF) (Burnett et al., 2009; Conrad et al., 2017), would have similar functional consequences in the normal activation mode of the HIV program, generating transcriptional fragility, and dampening the feedback loop as well as the magnitude of reactivation of a latent virus. Even when our experimental/mathematical modeling framework focused on NF-κB, which is a master regulator that becomes activated in response to many immune regulators, our discoveries are broadly applicable to different cell types, master regulators dictating proviral activation and immune cell signaling ligands and LRAs.

Weinberger et al. have shown Tat fluctuations could drive phenotypic diversity (Weinberger et al., 2005). Our results argue that this phenotypic diversity relies, at least partially, on the concept of “transcriptional fragility”. These observations indicate that the cellular state is a critical determinant for proviral transcription and escape from latency establishment. Thus, despite recent theories that the virus may function as a cell-autonomous unit (Razooky et al., 2015), the consensus in the field is that the status of the infected cell ultimately determines whether proviral transcription would be ‘on’ or ‘off’. In fact, it is believed that proviral transcription remains in the ‘off’ state without immune stimulation, which induces signaling events that generate a cellular state for the induction of host cell master regulators required for proviral transcription activation. This is because efficient formation of fully elongated and mature HIV transcripts requires sustained induction by the host cell activators, which will then promote *de novo* synthesis of Tat (Karn, 2011; Williams et al., 2007). Thus, importantly, without proper activation of the host phase (“fragility”), the magnitude of the viral phase, and consequently the Tat feedback loop, gets largely compromised (Dahabieh et al., 2015; Karn, 2011; Lassen et al., 2004; Mbonye and Karn, 2014; Ruelas and Greene, 2013).

Given KAP1 operates in primary T cells, it is possible that, as a consequence of the system’s “transcriptional fragility”, fluctuations in KAP1 protein levels in patient samples could affect the host phase, and ultimately impact the extent of the Tat feedback loop and the magnitude of HIV latency-reversal (thereby leading to proviral fate divergence). As such, single-cell heterogeneity in host phase responses could thus account for the large variations in latency-reversal observed both in different primary models and patient samples *ex vivo* (Spina et al., 2013). The system’s fragility is demonstrated in clonal CD4+ T cell lines where reduced KAP1 protein levels directly correlates with decreased, blunted proviral activation in response to immune stimulation. Thus, these results argue that stochasticity in the host phase (due to KAP1 level fluctuations in patients) could affect the feedback loop and impact on latency reversal potential. Despite mechanistic evidence in CD4+ T cell lines and primary cells, the proposed “fragility” model will have to be tested in patient-derived cells to provide *in vivo* relevance. To do so, future approaches would help overcome current technical difficulties to simultaneously measure KAP1 protein expression levels and HIV reactivation at the single-cell level.

Nonetheless, our discoveries have direct implications for HIV cure efforts in individuals who have full suppression of viral replication on ART. Resting memory CD4 T cells harboring stably integrated HIV genomes are capable of producing infectious virus upon T cell activation. Two current approaches for targeting the latent HIV reservoir from patients include ‘shock and kill’ (to reactivate all latent viruses and eliminate the cells containing induced virus through immune surveillance mechanisms) and ‘block and lock’ (to devise strategies for long-term, durable slicing of latent proviruses) (Sengupta and Siliciano, 2018). Even within one patient, the number of latent reservoirs and proviruses harboring replication-competent proviruses is immense and can lead to clonal expansion over time (Ho et al., 2013; Maldarelli et al., 2014; Wang et al., 2018). As such, several factors contribute to the pleiotropy in heterogeneous transcriptional responses by replication competent proviruses in patients including host circuit fragility and the integration landscape, among others. We argue that these latently-infected resting cells respond differently to T cell activation stimuli and LRAs as a consequence of the fragility of the transcriptional circuit in the host phase. Besides the possible fluctuations in activation of the viral phase (Tat function) previously observed (Weinberger et al., 2005), we argue the main component that must be taken into account during either of the proposed approaches to target the latent reservoir (Sengupta and Siliciano, 2018), is the level of activation of the host phase, which will set the threshold for activation of the cell-autonomous viral phase.

Intriguingly, beyond its proviral activating role, KAP1 appears to regulate other phases of the viral life cycle, such as integration (Allouch et al., 2011; Figueiredo and Hope, 2011). Thus, it seems reasonable to think it would be beneficial for the virus to integrate in sites of high chromatin accessibility bound by KAP1, which can then facilitate proviral activation (either in steady-state conditions or in response to immune signaling) thereby guaranteeing active replication. These possibilities open up a new area of investigation to test a potential KAP1 role in coupling both viral life cycle regulatory steps.

In addition to the KAP1 activating role in HIV proviral transcription, KAP1 has been previously linked to transcriptional repression of genes and retroelements in ES cells and other pluripotent states (Brattas et al., 2017; Ellis et al., 2007; Rowe et al., 2010; Rowe et al., 2013; Wolf and Goff, 2007, 2008, 2009). By interacting with unique members of the KRAB-ZnF family of DNA-binding factors, KAP1 mediates epigenetic silencing through deposition of repressive histone marks (H3K9me3) in ES cells (Wolf and Goff, 2009). It remains unclear, however, why KAP1 does not fulfill the expected repressive function in CD4+ T cell lines and in the primary T_CM_ model. It is possible the lack of KRAB-ZnF required for KAP1’s repressive role (Imbeault et al., 2017), and/or the nature of the cellular systems used (permissive/committed cell lineages, at difference to non-permissive backgrounds) both contribute to the lack of the well-established KAP1-mediated epigenetic silencing.

Interestingly, consistent with the previous theory, KAP1 could silence the expression of other viruses in a cellular system-dependent manner. Recent data by the Trono lab indicated that loss of KAP1 releases human CMV from latency in CD34+ hematopoietic stem cells (HSC), but not in permissive cells such as fibroblasts (Rauwel et al., 2015). Thus, KAP1 plays a dual role as repressor and activator depending on the cell-type and interacting protein complexes (Bunch et al., 2014; McNamara et al., 2016b) and could only confer the repressive phenotype in a context-dependent manner. It is worth noting that KAP1-mediated silencing of retroelements and genes during development is a mechanism that has been acquired during millions of years of evolution (Imbeault et al., 2017). As such, it makes sense that HIV (a relatively young virus in the evolutionary timescale) would not have undergone this suppressive regulatory mechanism by host cell factors like the KRAB-ZnF DNA-binding factors.

Finally, our findings provide a mechanistic explanation for the importance of the host phase of the HIV transcriptional program to ensure the virus is readily and robustly activated during infection to complete the pathogenic cycle. While HIV drives molecular innovation to fuel robust gene activation, it suffers from host phase fragility thereby influencing latent proviral transcription and homogeneous reactivation potential. Taken together, our discoveries have important implications for disease control and targeting in patients, and our experimental-mathematical modeling framework should provide a resource to guide the discovery of alternative HIV cure approaches.

## EXPERIMENTAL PROCEDURES

### Cell Culture

Jurkat CD4+ T cells (E6.1 clone, ATCC TIB152) and derivatives (Jurkat HIV E4, 10.6, 6.3, 8.4 and 9.2 clones, and Jurkat HIV *Tat-null* (2B2D clone) (Jordan et al., 2003; Pearson et al., 2008) (see **Table S1** for a complete list of cell lines used in this study) were maintained in RPMI 1640 media (HyClone, Cat. No. SH30027.FS) supplemented with 10% Fetal Bovine Serum (FBS) (Sigma, Cat. No. F4135) and 1X Penicillin/Streptomycin at 37°C with 5% CO_2_. The E4 and 2B2D clones derive from HIV-1 NL4-3 infectious molecular clone (Pearson et al., 2008) and 10.6, 6.3, 8.4 and 9.2 clones derive from the R7/3/GFP molecular clone and contain an *env* frame shift and GFP in place of *nef* (R7/E-/GFP) (Jordan et al., 2003). Jurkat cells stably expressing shRNAs were grown as above, but selected with the addition of 1 μg/ml of puromycin in case of the pLKO.1 cell lines or cell sorted (mCherry+), as indicated below, in the case of the pLVTHM cell lines. SupT1 cells (ATCC CRL-1942) were cultured in RPMI-1640 media containing 10% FBS and 2 mM L-Glu. U2OS (ATCC HTB96) and derivatives (shNT and shKAP1), HEK293T (ATCC CRL-11268) and HEK293FT (Thermo Fisher, Cat. No. 70007) cells were maintained in Dulbecco’s modified Eagle’s medium (DMEM) (HyClone, Cat. No. SH30022.FS) supplemented with 10% FBS and 1X Penicillin/Streptomycin (MP Biomedicals, Inc Cat. No. 091670049) at 37°C with 5% CO_2_. The Jurkat clones were treated with TNF-α (Sigma, Cat. No. T6674), SAHA (ApexBio, Cat. No. MK0683), Bryostatin (Sigma, Cat. No. B7431), Triptolide (Santa Cruz Biotechnologies, Cat. No. sc-200122) or Flavopiridol (Sigma, Cat. No. F3055) for the indicated time points and concentrations. Primary CD4+ T cells were isolated and cultured as indicated in the section “Generation of the Latency Model in Primary Central Memory T cells (T_CM_) and Analysis” below.

### Lentiviral Transduction and shRNA-mediated Knockdown

pLKO.1 non-targeting (NT, SHC002) and KAP1 (SHCLND-NM_005762) directed shRNA’s were obtained from Sigma. NT and KAP1 shRNAs were cloned into the ClaI and MluI restriction sites of the pLVTHM vector (**Table S2**) using standard molecular biology procedures (Ausubel et al., 1994). PLVTHM-expressing mCherry instead of GFP was previously described (Jadlowsky et al., 2014). The empty and NELF-E shRNA expressing pLVTHM vectors were kindly provided by J. Karn (Case Western Reserve University, Cleveland, OH) (Jadlowsky et al., 2014). The pLKO.1 and pLVTHM shRNA-containing vectors were transfected along with *gag/pol* [pSPAX vector] (Addgene) and VSV-G [pDM2 vector] (Addgene) into HEK293T cells for expression of competent lentiviruses. Cell supernatants were collected two days post-transfection. Viral transduction was done by spinoculation using 2×10^5^ cells, 8 µg/ml polybrene (Sigma, H9268), and unsupplemented RPMI-1640 to a final volume of 0.2 ml per well of a 96-well plate at room temperature for 2 hr at 400 g. Transduced cells were selected with puromycin (1 μg/ml) 2 days post-infection (for pLKO.1) or cell sorted on mCherry(+) cells (for pLVTHM). Cells were monitored for KD efficiency through standard western blot and RT-qPCR assays. For **Figure 5**, each cell line (Jurkat HIV Tat-shNT and Jurkat HIV Tat-shKAP1) was transduced with pTRIP-LUC (Schoggins et al., 2011) or pTRIP-Tat (see **Table S5**) in 96-well plates (2 plates per cell line at 0.2 ml lentiviral mix/well). Cells were then used in flow cytometry and RT-qPCR assays as indicated below. For **Figures S4A–C**, Jurkat HIV Tat-cell lines containing IPTG-inducible NT and KAP1 shRNAs (pLKO.1-puro-IPTG-3xLacO) (see **Table S2**) were created by transducing Jurkat HIV Tat-with the corresponding lentiviruses and selecting with puromycin (1 μg/ml) for 1 week. Selected cell lines were then treated for 2 days with three different IPTG concentrations (1, 10, and 100 μM) to precisely model the dosage effect of KAP1 protein levels on proviral transcription activation.

### Flow Cytometry Analysis

5×10^5^ cells per sample were transferred to an uncoated V-bottom 96-well plate (Nunc). The samples were spun down at 750 g for 5 min at room temperature and washed with 1X PBS (HyClone, Cat. No. SH30028.02). Washed cells were spun down again, and the 1X PBS was aspirated. Cells were fixed using 20 μl of 1% paraformaldehyde (PFA) (Sigma, Cat. No. P6148-500G) at room temperature for 10 min. The PFA was washed with 100 μl of PBS, spun down, buffer aspirated, and cell pellets resuspended in 100 μl of 1X PBS. A Stratedigm A600 HTAS 96-well plate reader (Stratedigm) was used to run the samples; lasers with a wavelength of 615 nm and 530 nm were used to measure mCherry and GFP, respectively. CellCapTure (Stratedigm) was used to visualize the running samples. 20,000 set count cells were analyzed per sample. Data analysis was performed with FloJo version 10.1 (TreeStar Inc). For CXCR4 detection, cells were stained with Fixable Viability Dye eFluor 450 (eBioscience, Cat. No. 65-0863-14) and anti-human CD184/CXCR4-PE (BD Biosciences, Cat. No. 555974) for 30 min. HIV-1_NL4-3_-infected cells were analyzed by flow cytometry first by staining with Fixable Viability Dye eFluor 450 (eBioscience, Cat. No. 65-0863-14) and anti-CD4-APC (clone S3.5, Invitrogen, Cat. No. MHCD0405) for 30 min. After washing with 1X PBS containing 3% FBS, cells were fixed and permeabilized using Cytofix/Cytoperm (BD, Cat. No. 554714) for 30 min and then stained with anti-p24-FITC (KC57, Beckman Coulter, Cat. No. 6604665). Flow cytometry was performed with a BD LRS Fortessa X-20 flow cytometer using FACSDiva acquisition software (Becton Dickinson, Mountain View, CA). Data was analyzed with Flow Jo version 10.1 (TreeStar Inc).

### Cell Sorting

One day post-transduction, cells were collected for sorting, washed with sterile 1X PBS, and resuspended in 10% RPMI-1640 media containing 10% FBS in 1X PBS. Cells were then transferred to 5 ml sterile polypropylene collection tubes (Falcon, Cat. No. 352063) containing 1 ml of 10% complete RPMI-1640 media in PBS, and analyzed directly or kept at 4°C until sorting (within 1 hr). A BD FACS Aria II (Becton Dickinson) was used (UTSW Flow Cytometry Core Facility) to sort live mCherry+/GFP-cells. A purity check was run after 1×10^6^ cells had been sorted. The cells were spun down and resuspended in 5 ml of complete RPMI-1640 media and grown in T-25 flasks (Corning, Cat. No. 430639) for use in western blots and RT-qPCR assays.

### RNA Extraction and RT-qPCR Assay

Isolation of total RNA was done using the Quick-RNA miniprep kit (Zymo, Cat. No. R1055). RNA quality was assessed by computing the RIN index (RNA Integrity Number) by running the samples on a 2200 Tapestation (Agilent) and was always RIN>9.5. First strand cDNA synthesis was done using M-MuLV Reverse Transcriptase (New England Biolabs, Cat. No. M0253) with oligo(dT)_18_ and random decamers (Thermo Fisher, Cat. No. AM5722G). Quantitative PCR was performed with a SybrGreen master mix on an ABI7500 instrument (Applied Biosystems). Ct values were obtained as previously described in detail (McNamara et al., 2013). The fold change of the target mRNA levels relative to control was calculated as 2^-ΔΔCt^. A list of DNA oligonucleotides used in RT-qPCR assays can be found in **Table S3**.

### ChIP-qPCR Assays

ChIP assays in Jurkat cells were performed as previously described (McNamara et al., 2016b). Purified cell nuclei were sonicated 60 cycles (30 sec on/30 sec off) on a Bioruptor UCD-300 water bath (Diagenode) to obtain DNA fragments of an average size of ∼300 bp. 5 μg of antibody were conjugated to 50 μl of 50% slurry protein G Dynabeads (Thermo Fisher, Cat. No. 10003) at 4°C for 2 hr and added to purified sonicated cell nuclei as follows: 1×10^7^ cell nuclei for Pol II, and 2.5×10^7^ cell nuclei for Cdk9, KAP1, and IgG (see **Table S4** for complete list of antibodies and their information). ChIP assays were performed with protein extracts from the indicated cells and using the antibodies indicated followed by qPCR with a series of amplicons mapping throughout the entire provirus mentioned at the top of the schematic to monitor factor interactions with the HIV genome.

The ChIP-qPCR data was normalized using the “Percent Input Method”, which includes normalization for background and Input chromatin used for each ChIP. ChIP signals were divided by signals obtained from the Input sample (1% of starting chromatin), which represents the amount of chromatin used in the ChIP. Values represent the percentage (%) of input DNA immunoprecipitated (IP DNA) presented after background (normal IgG) substraction, and are the average of three independent experiments.

### GAL4 Plasmid DNAs and Luciferase Assay

U2OS shNT and U2OS shKAP1 cell lines were seeded onto 48-well plates and transfected with a mix of DNAs (250 ng total DNA/well) and 0.5 μl Polyjet (SignaGen, Cat. No. SL100688) per well. For the experiment in **Figure 5A**, both cell lines were transfected with a pCDNA3.1-HIV-LTR-FFL reporter (25 ng/well) and increasing amounts of a pcDNA4/TO-Tat:Strep plasmid and carrier DNA (pBluescript II KS + Agilent, Genbank X52327) to complete 250 ng total DNA per well. For the experiment in **Figure 7**, the luciferase reporter plasmid is a pcDNA3.1+ vector (Thermo Fisher, Cat. No. V79020) containing a minimal LTR promoter plus 5xGAL4 binding sites and the activator plasmid containing yeast GAL4 DNA-binding domain alone or fused to the indicated P-TEFb subunit (CycT1, Cdk9, and Cdk9_T186A_ non-functional mutant) as previously described (D’Orso et al., 2012) (see **Table S5**). Firefly luciferase reporter activities were normalized to a constitutive CMV Renilla (RL) luciferase expressor using the dual luciferase kit (Promega, Cat. No. E1960). Luciferase of cell supernatants in SupT1 and primary CD4+ T cell infections was measured using Nano-Glo Luciferase Assay System (Promega, Cat. No. N1110).

### Western blot Assays

Total protein extracts from 0.2×10^6^ cells (∼20 μg, as quantitated using the Pierce BCA protein assay kit (ThermoFisher, Cat. No. 233225)) were electrophoresed on home-made 10% polyacrylamide SDS-PAGE gels using 1X Tris-Glycine-SDS running buffer prepared from a 10X stock (Bio-Rad, Cat. No. 1610732), and then transferred onto 0.45 μM nitrocellulose membranes (Bio-Rad, Cat. No. 1620115) using a standard Towbin transfer buffer (20% methanol, 25 mM Tris, 192 mM glycine, pH 8.3). Once transfer was complete, membranes were blocked in Tris-buffered saline (TBS) containing 0.2% Tween-20 (FisherBioreagents, Cat. No. BP337-500) and 5% non-fat dry milk (LabScientific, Cat. No. M0841) for 2 hr, and incubated with primary antibodies at 4°C from 1 hr to overnight. See list of all primary and secondary antibodies and their concentrations used in **Table S4**. Once the blotting was complete, membranes were incubated for 5 min with Clarity Western ECL substrate (Bio-Rad, Cat. No. 1705061) and exposed to film (Phenix Research Products, Cat. No. F-BX57 and F-BX810). Films were then scanned, cropped in Photoshop (Adobe) and directly used to make the figures in Illustrator (Adobe) without any further manipulation. When indicated (**Figure 7**), signal intensities in western blots were quantified using Image J (NIH).

### Transduction of Jurkat HIV Tat-Cell Lines for Flow Cytometry, RNA, Protein and ChIP Assays

pTRIP-LUC (Schoggins et al., 2011), pTRIP-Tat, and pTRIP-Tat C22G plasmid DNAs (**Table S5**) were transfected into HEK293T cells for lentiviral production as previously mentioned. The NT and KAP1 shRNA-expressing Jurkat HIV Tat-(2B2D) cell lines were transduced with the pTRIP lentiviruses indicated above. Cells were collected on day one, three, and five days post transduction for FACs analysis (GFP, RFP), or the indicated assays, as previously mentioned.

### Virus Production

For **Figure 3** and **Figure S2**, pseudotyped viruses (pNL4.3-delta*Env*-nLuc-2A*Nef*-VSVG) were produced by co-transfecting pNL4.3-delta*Env*-nLuc-2A*Nef* (containing NanoLuc (Promega)) (Martins et al., 2016) and pCMV-VSV-G (Addgene, Cat. No. 8454) (in a 2.5:1 plasmid DNA ratio) into HEK293T cells using calcium phosphate. After 2 days, cell supernatants were collected and filtered with a Millex-GP syringe filter unit, 0.22 μm, polyethersulfone, 33 mm, gamma sterilized (Millipore-Sigma, Cat. No. SLGP033RS). Viruses were tittered on SupT1 cells and stored at −80°C when needed. SupT1 cells were infected by viruses in a series of amounts. P24 expression was checked by flow cytometry 2 days post-infection.

### Generation of CRISPR-Cas9 Knockout on HIV Latency Models

For **Figure 3** and **Figure S2**, cells were infected with pseudotyped viruses (pNL4.3-delta*Env*-nLuc-2A*Nef*-VSVG) for 2 days. After amplification as indicated in the Figure, CD4+ cells were isolated using Dynabeads CD4 Positive Isolation Kit (Thermo Fisher, Cat. No. 11331D). TracrRNA (IDT, Cat. No. 1072532) and guide RNAs (IDT, scrambled gRNA (Cat. No. 1072544); CXCR4 gRNA: 5’-GAAGCGTGATGACAAAGAGG-3’; NF-κB p65 subunit gRNA: 5’-GAGGGGGAACAGTTCTGAAA-3’; KAP1 gRNA: 5’-ACGTTCACCATCCCGAGACT-3’) were mixed and heated at 95°C for 5 min. Cas9 (IDT, Cat. No. 1081058) was added to the RNA mixture [3.23 μg Cas9 protein and 21.6 pmol gRNA] and incubated for 20 min. CD4+ cells were washed with PBS and resuspended in 10 μl of Buffer R (for SupT1) or Buffer T (for primary CD4+ T cells) of Neon Transfection System kit (Thermo Fisher, Cat. No. MPK1096). Pre-assembled Cas9-gRNA ribonucleoprotein (RNP) complexes were electroporated into cells using Neon Transfection System (Thermo Fisher, Cat. No. MPK5000). After 2 days, cell viability and CXCR4 staining were performed. Cells were then seeded into 96-well plates, treated with 10 ng/ml PMA (Sigma, Cat. No. P8139) or vehicle (DMSO 99.7%, ACROS Organics, Cat. No. 610420010) for 2 days. Luciferase and intracellular p24 levels were recorded by luciferase assays and flow cytometry, respectively, as indicated above.

### Generation of the Latency Model in Primary Central Memory T cells (T_CM_) and Analysis

Peripheral blood mononuclear cells (PBMCs) were isolated from healthy donors. Naïve CD4+ T cells were isolated and T_CM_ cells were generated and infected as previously described (Bosque and Planelles, 2009, 2011; Martins et al., 2016; Messi et al., 2003). Briefly, naïve CD4+ T cells were obtained by magnetic isolation (Stemcell Technologies, Cat. No. 19555) from healthy donor blood samples (Gulf Coast Regional Blood Center). Naïve CD4+ T cells were activated using human anti-CD3/CD28-coated magnetic beads (Thermo Fisher, Cat. No. 11161D) in the presence of anti-IL4 (2 µg/10^6^ cells, Peprotech, Cat. No. 500-P24), anti-IL12 (4 µg/10^6^ cells, Peprotech, Cat. No. 500-P154G) and tumor growth factor (TGF-β1) (0.8 µg/10^6^ cells, Peprotech, Cat. No. 100-21) for 3 days. After 3 days, magnetic beads were removed, cells were washed and maintained at a concentration of 10^6^ cells/ml in media containing 30 IU of human IL-2 (Roche, Cat. No. 126). HIV-1_NL4-3_ virus was generated in HEK293FT cells using calcium phosphate. T_CM_ cells were then infected with HIV-1_NL4-3_ by spinoculation at a multiplicity of infection (MOI) of 0.6 using a concentration of 2×10^6^ cells/1 ml and centrifuged for 2 hr at 37°C and 162 g. Following infection, cells were cultured in 96-well round bottom plates (10^5^ cells/100 µl/well) for 3 days (from day 7 to 10). At day 10, cells were cultured in standard tissue culture flasks at cell density of 10^6^ cells/ml. At day 13, 1 µM of nelfinavir (NIH AIDS Reagent Program, Cat. No. 4621) was added to the cells for viral suppression.

### Crosslinking of Primary T_CM_ Cells for ChIP Assays

T_CM_ cells (∼2×10^7^) were pelleted by centrifugation (600 g for 5 min at room temperature) and resuspended at a density of 1×10^7^ cells/ml in 0.5% methanol-free formaldehyde (Thermo Fisher Cat. No. 28908) diluted in 1X PBS. Cells were nutated for 5 min at room temperature. Glycine (0.15 M) was added to quench crosslinking and cells nutated for 10 min at room temperature. Cells were then pelleted at 750 g for 5 min at 4°C and 2X with cold PBS. Snap-frozen cell pellets were kept at - 80°C until sonication as indicated above. Briefly, T_CM_ cells were processed for ChIP assays like Jurkat cells. 2×10^7^ cell nuclei were used per ChIP assay with IgG, Pol II, CDK9, and KAP1 antibodies as indicated in **Figure 3**.

### Statistical analyses

Student’s *t*-test was used to determine statistical significance when indicated. We considered *P <* 0.05 to be statistically significant.

### Mathematical Modeling

See Supplementary Information for a complete description of the model.

Refer to the **Supplemental Experimental Procedures** for additional information.

## SUPPLEMENTAL INFORMATION

Supplemental information includes 4 figures, 7 tables, and can be found appended to this manuscript.

## AUTHOR CONTRIBUTIONS

E.L.M., C.V.F and I.D. conceived the study. A.B.S., Y.Z., V.P. and I.D. developed the experimental outline for the primary cell model. N.G.R. prepared DNA clones used in this study and assisted E.L.M in the lentiviral assays and flow cytometry analysis. E.L.M., A.B.S., Y.Z., and I.D. conducted mechanistic studies. A.B.S, E.L.M, I.D. and Y.Z. conducted primary cell studies. C.V.F. developed the mathematical models, performed predictive simulations and analyzed the outcome in the context of the experimental data. E.L.M., C.V.F., V.P. and I.D. wrote the manuscript.

## ACKNOWLEDGEMENTS

We thank the UT Southwestern Flow Cytometry Core for its assistance with flow cytometry and cell sorting. We are grateful to J. Karn for generously sharing the Jurkat HIV E4 and 2B2D clones. The 10.6, 6.3, 8.4, and 9.2 Jurkat HIV clones were obtained from the NIH AIDS Reagent Program kindly deposited by E. Verdin. Research reported in this publication was supported by the National Institute of Allergy and Infectious Diseases of the NIH under award numbers R01AI114362 and R33AI116222, and Welch Foundation grant number I-1782 (to I.D.), by the National Institute of Allergy and Infectious Diseases of the NIH under award numbers R21 AI123035-01 and R33 AI122377 (to V.P.), and by the National Institute of Allergy and Infectious Diseases of the NIH under award numbers U01AI111598 (to C.V.F.). N-G.P.R was supported by the NIH Pharmacological Sciences Training grant number GM007062 and a 2016 pre-doctoral fellowship from the Ford Foundation.

