## Supplemental Information for "Transcriptional Circuit Fragility Influences HIV Proviral Fate"

Supplemental Data: pages 2–10

Extended Experimental Procedures: pages 11–19

Supplemental Tables: pages 20–26

Supplemental References: page 27

**Supplemental Data**

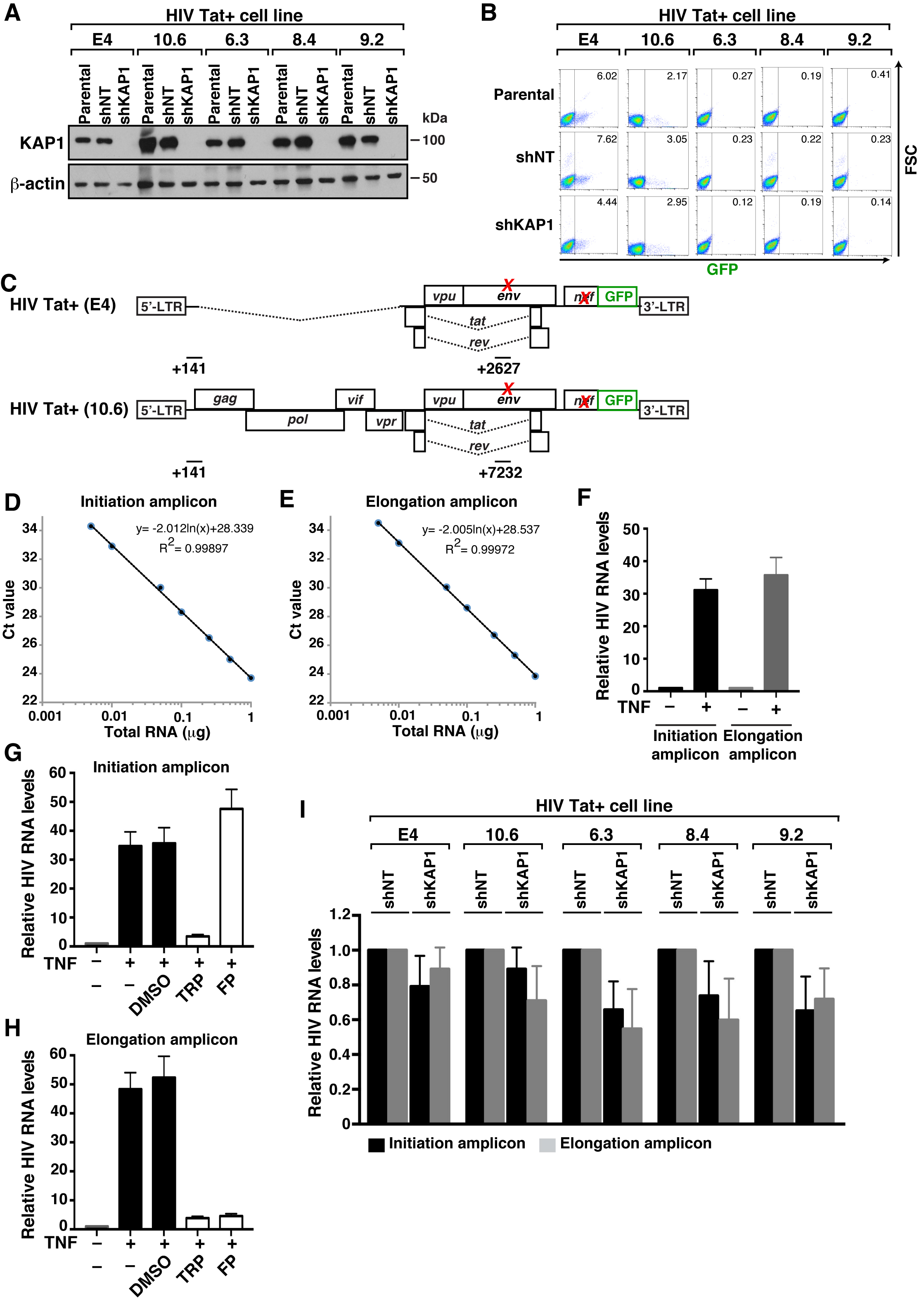

**Figure S1. Loss of KAP1 Does not Appear to Affect Basal HIV Transcription. Related to Figure 2**

(A) Western blots of the different HIV Tat+ cell-based models created as described in **Figure 2**.

(B) Flow cytometry analysis of the indicated cell-based models from panel A. FSC, forward scatter. The number in the quadrant denotes the average percentage of GFP+ cells (n = 3).

(C) Scheme of the HIV proviruses with the position of the amplicons used in RT-qPCR assays. The top scheme corresponds to the HIV Tat+ (E4) provirus, while the bottom scheme corresponds to HIV Tat+ (10.6, 6.3, 8.4, and 9.2). The position of the initiation amplicon (+141) is indicated. The position of the elongation amplicon in the E4 proviral genome is +2627 respective to the TSS, and +7232 in the other proviruses because they contain full-length genomes.

(D-E) Standard curves for RT-qPCR assay. Total RNA from the HIV Tat+ (E4) clone treated with 25 ng/ml TNF for 16 hrs was serially diluted and seven aliquots between 0.005 ng and 1 μg were converted to cDNA using individual RT reactions before performing qPCR with the initiation amplicon (+141) and elongation amplicon (+2627). While the initiation amplicon only measures short, promoter-proximal transcripts, the elongation amplicon measures promoter-distal transcripts. PCR amplifications were performed in 20 μl reaction mixtures containing 10 μl of SYBR green master mix (Applied Biosystems), primers and 2 μl of cDNA. The plot demonstrates linear reverse transcription for the concentrations of RNA tested without any effect of RNA input beyond RT capacity. In this situation, both HIV short and long target transcripts (as well as the internal control *ACTB* (data not shown)) have linear RT efficiencies across all starting concentrations of RNA tested. The qPCR plots show threshold Ct values (y-axis) as a function of increasing RNA concentrations (x-axis).

(F) Relative HIV RNA levels of HIV Tat+ (E4) treated (+) or not (-) with 25 ng/ml TNF for 2 hr by RT-qPCR and normalized to *ACTB* (mean ± SEM; n = 3).

(G) Quantification of short transcripts with the initiation amplicon (+141) from total RNA from the HIV Tat+ (E4) clone isolated after treatment with 25 ng/ml TNF alone for 2 hr or pre-treated with TRP, FP or DMSO for 30 min before the addition of TNF. Relative HIV RNA levels were normalized to *ACTB* (mean ± SEM; n = 3).

(H) Quantification of long transcripts with the elongation amplicon (+2627) from total RNA from the HIV Tat+ (E4) clone isolated after treatment with 25 ng/ml TNF alone for 2 hr or pre-treated with TRP, FP or DMSO for 30 min before the addition of TNF. Relative HIV RNA levels were normalized to *ACTB* (mean ± SEM; n = 3).

(I) Relative HIV RNA levels: initiation (black bars) and elongation (grey bars) transcripts quantified by RT-qPCR and normalized to *ACTB* (mean ± SEM; n = 3). Statistical significance was determined using unpaired Student’s *t*-test. **P* < 0.05, ***P* < 0.005, ****P* < 0.0005.

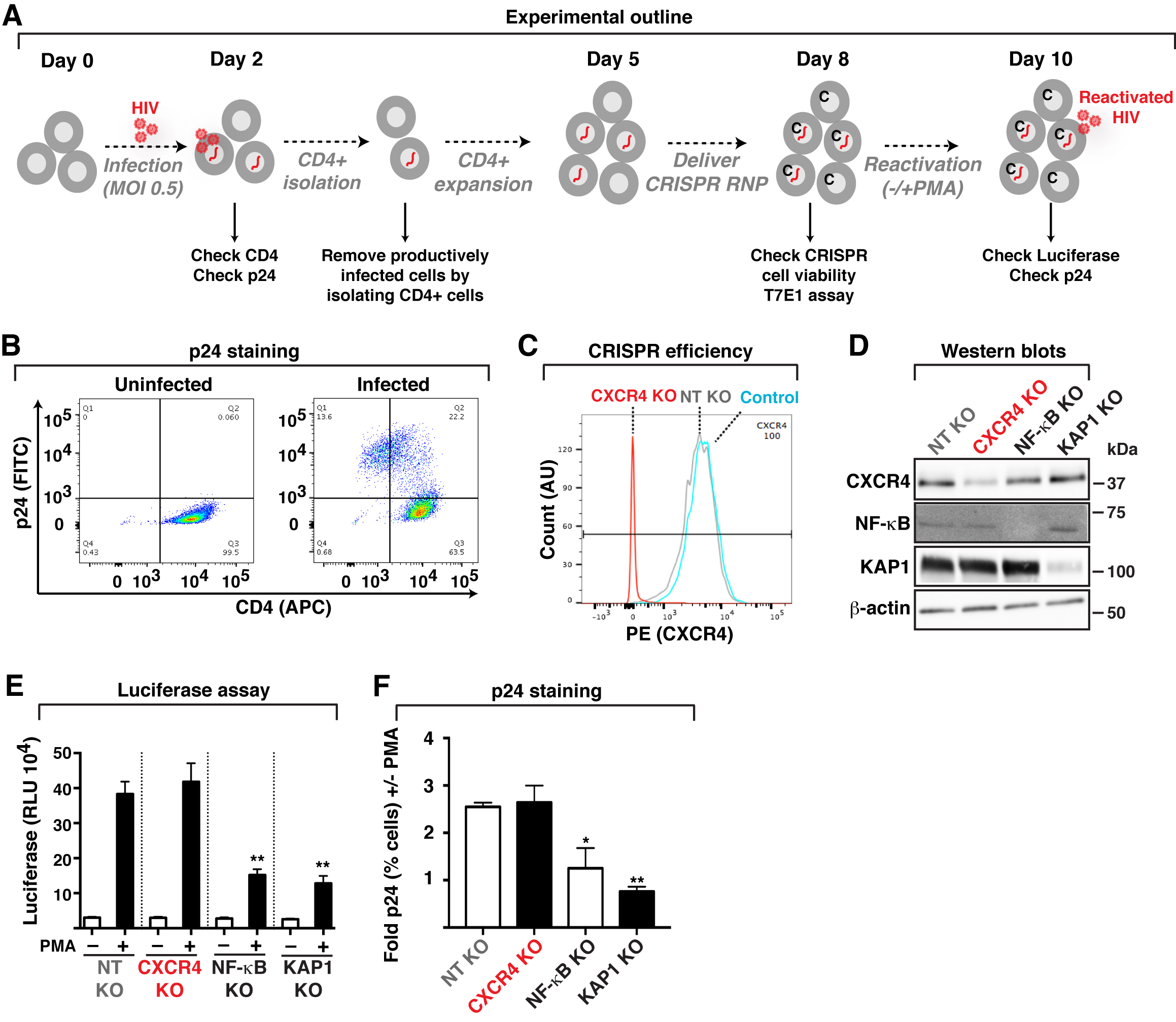

**Figure S2. KAP1 is Required for Latent HIV Reactivation in Response to Immune Signaling in Cell-based Models. Related to Figure 3**

(A) Experimental outline through which CD4+ SupT1 cells were used for infections with replication competent, pseudotyped HIV (pNL4.3-delta*Env*-nLuc-2A*Nef*-VSVG), and then used for CRISPR-Cas9–mediated knockout (KO) of CXCR4, NF-κB (p65 subunit) and KAP1, followed by reactivation assays. RNP, Cas9-gRNA RiboNucleoProtein complex. C, indicates cells containing the Cas9-gRNA RNP complex.

(B) FACS plots (CD4, HIV p24) of mock infected (uninfected) and HIV-infected cells as in panel (A).

(C) FACS plots (CXCR4) in control SupT1 cells (not nucleofected) and SupT1 nucleofected with Cas9-gRNA complexes for targeting CXCR4 and a non-target (NT) negative control.

(D) Western blots of SupT1 cells containing KO of specific host cell factors generated as in panel (A) with the indicated antibodies.

(E) Luciferase assay of SupT1 cells containing KO of specific host cell factors generated as in panel (A) and treated with PMA or vehicle (DMSO). Luciferase is expressed as relative luciferase units (RLU).

(F) p24 staining of SupT1 cells containing KO of specific host cell factors generated as in panel (A) and treated with PMA or vehicle (DMSO). The fold change in p24 staining (+/- PMA) is indicated. Statistical significance in panels (E) and (F) was determined using unpaired Student’s *t*-test. **P* < 0.05, ***P* < 0.005, ****P* < 0.0005.

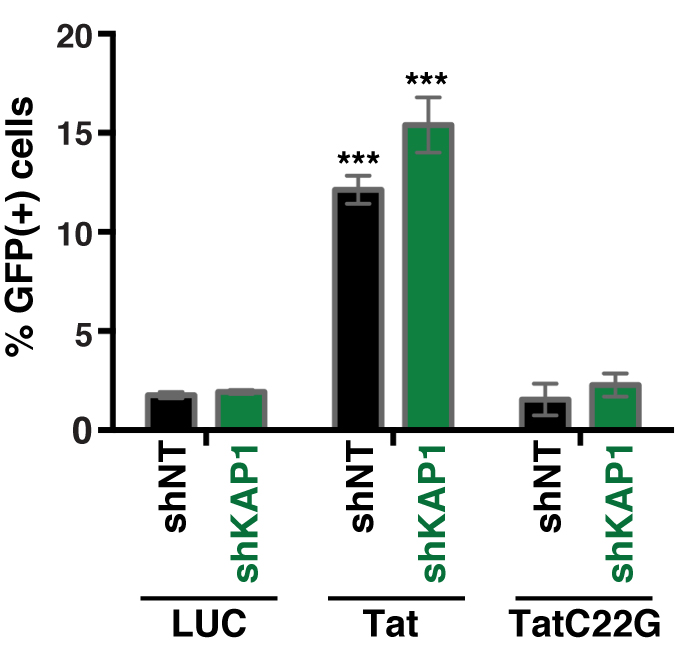

**Figure S3. Tat, but not a non-functional mutant, reactivates a latent HIV Tat- provirus. Related to Figure 5**

Quantification of GFP+ cells (percentage) in the Jurkat HIV Tat- shNT and shKAP1 cell lines transduced with pTRIP lentiviruses (5 ng p24) expressing firefly luciferase (LUC) as negative control, wild-type Tat or the C22G non-functional mutant. Statistical significance between Tat or TatC22G and LUC samples was determined using unpaired Student’s *t*-test. **P* < 0.05, ***P* < 0.005, ****P* < 0.0005.

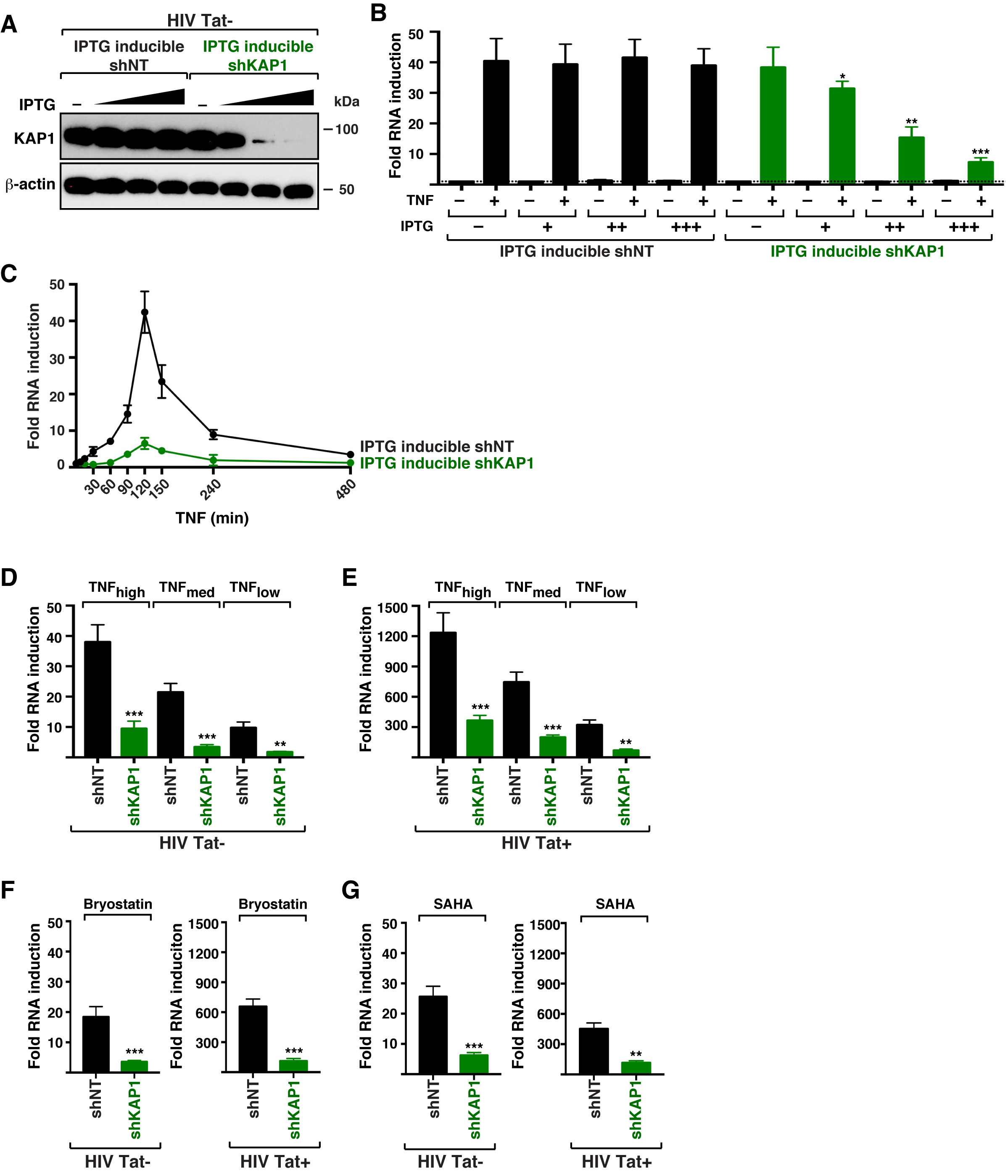

**Figure S4. Effects of KAP1 protein levels and graded, diverse immune stimuli on activation of the host and viral phases. Related to Figure 7**

(A) Western blots of the HIV Tat- (2B2D) IPTG-inducible shNT and shKAP1 cell lines untreated (-) or treated with three IPTG concentrations (1, 10, and 100 μM) for 2 days.

(B) Fold HIV RNA expression levels (+/- TNF) of the HIV Tat- (2B2D) IPTG-inducible shNT and shKAP1 cell lines from panel (A) untreated (-) or treated with three IPTG concentrations (1, 10, and 100 μM; +, ++, +++, respectively) for 2 days in the absence (-) and presence (+) of TNF stimulation for 2 hr and measured by RT-qPCR using the elongation amplicon (+2627) and normalized to *ACTB* (mean ± SEM; n = 3).

(C) Fold HIV RNA expression levels (+/- TNF) of the HIV Tat- (2B2D) IPTG-inducible shNT and shKAP1 cell lines from panel (A) treated with IPTG (100 μM) for 2 days in response to a time course TNF treatment and measured by RT-qPCR using the elongation amplicon (+2627) and normalized to *ACTB* (mean ± SEM; n = 3).

(D) Fold HIV RNA expression levels (+/- TNF) of HIV Tat- (2B2D) shNT and shKAP1 cell lines in the absence (-) and presence (+) of different amounts of TNF stimulation [TNF_high_ (25 ng/ml), TNF_medium_ (5 ng/ml), TNF_low_ (1 ng/ml)] for 2 hr and measured by RT-qPCR using the elongation amplicon (+2627) and normalized to *ACTB* (mean ± SEM; n = 3).

(E) Fold HIV RNA expression levels (+/- TNF) of HIV Tat+ (E4) shNT and shKAP1 cell lines in the absence (-) and presence (+) of different amounts of TNF stimulation [TNF_high_ (25 ng/ml), TNF_medium_ (5 ng/ml), TNF_low_ (1 ng/ml)] for 16 hr and measured by RT-qPCR using the elongation amplicon (+2627) and normalized to *ACTB* (mean ± SEM; n = 3).

(F) Fold HIV RNA expression levels (+/- TNF) of HIV Tat- (2B2D) and HIV Tat+ (E4) shNT and shKAP1 cell lines in the absence (-) and presence (+) of Bryostatin (10 nM) stimulation for 2 hr (HIV Tat-) or 16 hr (HIV Tat+) and measured by RT-qPCR using the elongation amplicon (+2627) and normalized to *ACTB* (mean ± SEM; n = 3).

(G) Fold HIV RNA expression levels (+/- TNF) of HIV Tat- (2B2D) and HIV Tat+ (E4) shNT and shKAP1 cell lines in the absence (-) and presence (+) of SAHA (500 nM) stimulation for 2 hr (HIV Tat-) or 16 hr (HIV Tat+) and measured by RT-qPCR using the elongation amplicon (+2627) and normalized to *ACTB* (mean ± SEM; n = 3). Statistical significance in panels (A-D) was determined using unpaired Student’s *t*-test. **P* < 0.05, ***P* < 0.005, ****P* < 0.0005.

**EXTENDED EXPERIMENTAL PROCEDURES**

**Introduction**

We seek to understand the functional interplay between host cell factors such as KAP1 and the cellular (NF-κB) and viral (Tat) transcriptional activators during HIV RNA synthesis and latency-reversal in response to immune stimulation. Notably, KAP1 allows for the initial NF-κB–mediated transcriptional “boost”, which facilitates robust Tat positive feedback loop. Conversely, loss of KAP1 blunts the initial “boost” thereby dampening Tat function and latency-reversal. Although the viral-driven phase of the transcriptional program is “minimalist” (because of the bypass of host cell requirements), the strict dependence of cellular factors for the host phase makes the complete circuit “fragile”, thus revealing key information that must be contemplated for HIV cure strategies.

**Model Overview**

We are considering four components to model HIV RNA synthesis by the combined action of the host and viral phases. In cells expressing normal KAP1 levels (i) with and (ii) without feedback loop, and in cells where KAP1 expression is lost (iii) with and (iv) without feedback loop. In normal conditions (cells expressing normal KAP1 levels), one can distinguish between an early KAP1-dependent “boost” and a later KAP1-independent phase of HIV RNA synthesis.

**Step (0)**: Involving multiple feedback loops, NF-κB translocates from the cytosol to the nucleus after binding of TNF to its receptor and following activation of the IκB kinase (IKK).

**Step (1)**: RNA synthesis is initiated by NF-κB binding to the viral promoter (*k_on_*), which uses KAP1/CDK9 as co-activator for HIV proviral transcription activation.

**Step (2)**: Once at the viral promoter, KAP1 promotes CDK9 delivery and activation (*k_act(h)_*). However, KAP1/CDK9 activity decays quickly (*k_deact(h)_*) as a consequence of NF-κB dissociation from the promoter (*k_off_*).

**Step (3)**: This initial “boost” promotes the synthesis of HIV transcripts (*k_synth(h)_*), which can be degraded (*k_decay_*).

**Step (4)**: Alternatively, HIV RNAs serve as templates for translation (*k_trans_*) leading to Tat synthesis.

**Step (5)**: Although NF-κB concentration at the promoter decreases quickly as a consequence of its dissociation from the template DNA (*k_off_*), Tat itself takes the place of KAP1 by recruitment and activation/deactivation of CDK9 in the viral phase with kinetic activation and deactivation parameters *k_act(v)_* and *k_deact(v)_*, respectively.

**Step (6)**: As a consequence of Tat activity (Step (5)), the positive feedback loop (*k_fb_*) becomes dominant leading to robust and sustained HIV RNA synthesis (*k_synth(v)_*).

Without feedback loop, HIV RNA synthesis receives an initial KAP1-dependent “boost” by NF-κB. However, with NF-κB diminishing as a consequence of its dissociation from the promoter (*k_off_*) and re-translocation to the cytoplasm, HIV transcription soon returns to the low steady-state level of the basal transcription rate; and as a consequence, the feedback loop (*k_fb_*) does not operate normally. Loss of KAP1 virtually abolished the initial transcriptional “boost” mediated by NF-κB; consequently, the feedback loop is largely reduced in magnitude compared with the KAP1 positive scenario. Furthermore, without feedback loop, the levels of HIV RNA synthesis remain extremely low and indistinguishable from basal activity.

**Assumptions**

(1) Basal HIV RNA synthesis remains constant over time and is very low compared to RNA synthesis induced in response to NF-κB and Tat activation (thereby becoming depreciable).
(2) Basal HIV RNA does not contribute to the pool of molecules that generate fully mature HIV RNAs leading to viral products to perpetuate the infection.
(3) The overall effect of NF-κB activation involves the canonical positive/negative feedback loop due to NF-κB binding to the promoter (leading to activation) followed by its release (leading to deactivation).
(4) NF-κB–promoter association and dissociation is induced after activation by TNF.
(5) KAP1 is already bound to CDK9 and dissociates from the promoter with the given rate.
(6) Tat translation explicitly requires NF-κB–mediated HIV RNA synthesis in response to immune stimulation.
(7) Molecular processes involved in the Tat positive feedback loop are non-limiting and can be reduced to overall rates.

**Variables**

Time is simulated in discrete steps according to the simulation implementations used (deterministic or stochastic). [*RNA*](*t*), [*Tat*](*t*), [*NF-κB*](*t*), [*KAP1*](*t*) and [*TNF*](*t*) describe the amount of HIV RNA, Tat, NF-κB, KAP1 (in the nucleus) and TNF, respectively.

**Parameters**

See **Table S6**.

(1) *k_on_* and *k_off_* represent the spontaneous association and dissociation of transcription factor and co-activators to the proviral promoter, respectively.
(2) *k_act(h)_*, *k_deact(h)_*, describe the rates of KAP1-mediated recruitment and activation/deactivation of CDK9 in the host phase, respectively.
(3) *k_synth(h)_* describe the rate of HIV RNA synthesis by NF-κB (initial transcriptional boost; host phase).
(4) *k_act(v)_*, *k_deact(v)_*, describe the corresponding rates of Tat–CDK9 association and dissociation from the promoter, respectively.
(5) *k_synth(v)_* describe the rate of HIV RNA synthesis by Tat (feedback loop; viral phase).
(6) *k_trans_* and *k_decay_* describe the rate of translation and decay of Tat, respectively.
(7) *k_fb_* represents active Tat function in a CDK9-dependent fashion.

**Physical Basis for Parameters**

We assume a well-mixed scenario for our simulations. Kinetic parameters have been either taken from published data together with the underlying experimental conditions, or have been fitted using measured RNA concentrations and JSim's Simplex non-linear steepest-descent algorithm (Butterworth et al., 2013).

**Ordinary Differential Equations (ODE)**

**Table S7** shows a deterministic approximation of the model as system of six ODEs and a seventh equation of the total concentration of RNA in the system. We used a set of different initial conditions for KAP1 to assess its effect on the dynamics of the model. We consider basal and stimulated RNA expression separately. The dynamics of the non-basal RNA is dependent on activation by NF-κB, KAP1 and Tat. The dynamics of stimulated RNA synthesis in our dynamic model (**Table S7**) is described by ODE (1):

$$\frac{d\left[ RNA \right]}{dt}=\mu_{RNA}\frac{\left[ NF\kappa B \right]}{k_{Mm}+\left[ NF\kappa B \right]}+\tau_{RNA}\left[ KAP1 \right]\left[ NF\kappa B \right]+k_{synth(h)}\left[ KAP1 \right]\left[ Tat \right]+k_{synth(v)}\left[ Tat \right]-k_{decay}\left[ RNA \right]$$

(1)

The first term describes an overall, Michaelis-Menten type, dynamics regulated by NF-κB, with $\mu_{RNA}=k_{\text{cat}\cdot}\left[ NF\kappa B \right]_{0}$ describing the maximal reaction velocity with $\left[ NF\kappa B \right]_{0}$ denoting the initial concentration of NF-κB, and $k_{Mm}=\left( k_{\text{off}}+k_{\text{cat}\cdot} \right)/{k_{\text{on}}}$ being the corresponding Michaelis-Menten constant.

The second term denotes the additional effect of KAP1 on transcription together with NF-κB. Overall activation rate is $\tau_{RNA}$ with the implicit inclusion of activation rate $k_{act(h)}$ (see below). The third and fourth terms refer to KAP1-initiated and KAP1-independent; respectively, Tat translation, and further contribution by Tat through the feedback loop. Given the stimulated RNA, which is further translated into Tat, and its concentration $\left[ RNA \right],$ the dynamic of Tat translation with [*Tat*] denoting the concentration of expressed Tat protein is then given by the following equation:

$$\frac{d\left[ Tat \right]}{dt}=k_{trans}\left[ RNA \right]-d_{Tat}\left[ Tat \right]$$

Tat is known to establish a positive feedback loop via binding to the TAR RNA stem-loop formed at the 5’-end of nascent viral pre-mRNAs. Following (Razooky and Weinberger, 2011), we ‘lump’ many of the detailed molecular interactions known to take place in this process into two parameters to generate a minimal model of HIV provirus *trans*-activation. The resulting minimal model can be described by ODE (2):

$$\frac{d\left[ Tat \right]}{dt}=k_{trans}\left[ RNA \right]+\mu_{Tat}\frac{\left[ Tat \right]}{k_{MTat}+\left[ Tat \right]}-d_{Tat}\left[ Tat \right]$$

(2)

We use the same terms as in Equations (1) and (2); however, the middle term represents a saturable positive-feedback loop, where $\mu_{Tat}$ represents the positive-feedback strength, and $k_{MTat}$ is the saturation constant of the system. Similar to the Michaelis-Menten approach we employed for the NF-κB–regulated dynamics in Equation (1), we use a Michaelis-Menten dynamics to model the Tat feedback loop in Equation (2). The corresponding maximal reaction velocity (*μTat*) is described by *μTat = k_fb_ [Tat]_0_ and k_MTat_ = (k_deact(v)_ + k_fb_)/k_act(v)_ being the* Michaelis-Menten constant. *d_Tat_* refers to the degradation rate of Tat.

Other reactions included in the model are the activation of NF-κB by TNF, transport of KAP1 to/from the nucleus and TNF signaling. We simplified the complex feedback loop between TNF, the inhibitor of NF-κB kinase (IKK), other kinases (such as TAK1), *IκBα*, A20 and *NF-κB* into a linear module capturing the overall dynamics of NF-κB activation by TNF.

$$\frac{d\left[ NF\kappa B \right]}{dt}=\beta\left[ TNF \right]-d_{NF\kappa B}\left[ NF\kappa B \right]$$

(3)

Kinetic parameter *β* describes the overall NF-κB activation (including translocation to the nucleus), whereas $d_{NF\kappa B}$ denotes the deactivation rate.

Similar to the simplification of the TNF ↔ NF-κB feedback loop, we streamline the translocation of KAP1 from the nucleoplasm to the promoter/chromatin, activation/deactivation of the promoter by KAP1, involving CDK9, and KAP1 relocation to the nucleoplasm by overall reactions with the implicit activation rate in the host phase $k_{act(h)}$ (see explanation to Equation (1)) and deactivation rate in the host phase $k_{deact(h)}$:

$$\frac{d\left[ KAP1 \right]}{dt}=\rho-d_{KAP1}\left[ KAP1 \right]$$

(4)

with import rate *ρ* and export/deactivation rate $d_{KAP1}$ (implicitly including $k_{deact(h)}$), yielding a stationary state concentration in the nucleus of $\left[ KAP1 \right]=\frac{\rho}{d_{KAP1}}$.

The activation by TNF is modeled by a simple exponential decay from a finite value at *t* = 0 with decay rate $d_{TNF}$.

$$\frac{d\left[ TNF \right]}{dt}=-d_{TNF}\left[ TNF \right]$$

(5)

The differential equation that describes the basal expression is:

$$\frac{d\left[ RNAbasal \right]}{dt}=\alpha-d_{RNAbasal}\left[ RNAbasal \right]$$

(6)

yielding a steady state of $\left[ RNAbasal \right]=\frac{\alpha}{d_{RNAbasal}}.$ *dRNA_basal_* denotes the degradations rate of basal RNA.

Together with the non-basal RNA, the total, measured RNA is then:

$$\left[ RNAtot \right]=\left[ RNA \right]+[RNAbasal]$$

(7)

**Stochastic Description**

RNA, Tat, KAP1, TNF, Pol II and NF-κB are actually simulated as discrete molecules in a well stirred mixture in which stochastic Poisson processes act. Such Poisson processes are well described by the Chemical Master Equation (Hahl and Kremling, 2016) capturing the corresponding reaction system.

$$KAP1\text{ initiation}: KAP1+NF\kappa B\underset{\to}{\tau_{RNA}}KAP1+NF\kappa B+RNA$$

(8)

$$NF\kappa B \text{initiation}: \left( \text{DNA/Pol}\text{ II} \right)+NF\kappa B\begin{matrix} k_{off} \\ \leftrightharpoons\\ k_{on} \end{matrix}\left( \text{DNA/Pol II} \right)\cdot NF\kappa B\underset{\to}{k_{cat}}NF\kappa B+RNA$$

(9)

$$Tat\text{ induced (}KAP1\text{ dependent) transcription}: KAP1+Tat\underset{\to}{k_{synth(h)}}KAP1+Tat+RNA$$

(10)

$$Tat\text{ induced (}KAP1\text{ independent) transcription}: Tat\underset{\to}{k_{synth(v)}}Tat+RNA$$

(11)

$$RNA\text{ degradation}: RNA\underset{\to}{k_{decay}}\emptyset$$

(12)

$$Tat\text{ translation}: RNA\underset{\to}{k_{trans}}Tat+RNA$$

(13)

$$Tat \text{feedback}: \left( RNA \right)+Tat\begin{matrix} k_{deact} \\ \leftrightharpoons\\ k_{act} \end{matrix}\left( RNA \right)\cdot Tat\underset{\to}{k_{fb}}2 Tat$$

(14)

$$Tat \text{degradation}: Tat\underset{\to}{d_{Tat}}\emptyset$$

(15)

$$NF\kappa B \text{activation by }TNF: TNF\underset{\to}{\beta}TNF+NF\kappa B$$

(16)

$$NF\kappa B \text{deactivation}: NF\kappa B \underset{\to}{d_{NF\kappa B}}\emptyset$$

(17)

$$KAP1 \text{"transport from cytosol to nucleus"}: \emptyset\underset{\to}{\rho}KAP1$$

(18)

$$KAP1/CDK9 \text{activity decay and relocalization}: KAP1\underset{\to}{d_{KAP1}}\emptyset$$

(19)

$$TNF \text{deactivation}: TNF \underset{\to}{d_{TNF}}\emptyset$$

(20)

KAP1 initiation denotes the contribution of KAP1 to the host phase and NF-κB initiation denotes activation of the host phase.

**Simulations**

Both the deterministic as well as the stochastic simulation uses standard procedures according to the implementation of the corresponding simulation software (see section “Implementation” below). In the case of the stochastic simulations, 100 trajectories for each run have been calculated.

**Implementation**

We implemented the deterministic simulation of the corresponding ODEs [equations (1) – (7)] within JSim v2.15 in JSim’s own Mathematical Modeling Language (MML). The Dormand-Prince explicit Runge-Kutta method of order 5(4) for non-stiff equations (Dopri5) was used for simulation, with a fallback option to the implicit Runge-Kutta method of variable order (Radau; solver setting to “auto”). Parameter optimization for unknown parameters was performed using the simplex method. The stochastic version of the dynamic model was implemented in StochSS (Drawert et al., 2016) [see equations (8) – (20)]. StochSS provides implementations of several exact stochastic simulation algorithms (SSA), including the direct method, optimized direct method and composition-rejection method. These methods all generate exact samples (trajectories) from the chemical master equation. After model analysis, StochSS automatically chooses the appropriate algorithm.

**Table S1. Cell lines used and created in this study**

| **Cell line** | **Laboratory** | **Reference** |
| --- | --- | --- |
| Jurkat E4 | Jonathan Karn | (Pearson et al., 2008) |
| Jurkat E4 NT shRNA | Iván D’Orso | Created in this study |
| Jurkat E4 KAP1 shRNA | Iván D’Orso | Created in this study |
| Jurkat E4 NELF-E shRNA | Iván D’Orso | Created in this study |
| Jurkat 10.6 | Eric Verdin | (Jordan et al., 2003) |
| Jurkat 10.6 NT shRNA | Iván D’Orso | Created in this study |
| Jurkat 10.6 KAP1 shRNA | Iván D’Orso | Created in this study |
| Jurkat 10.6 NELF-E shRNA | Iván D’Orso | Created in this study |
| Jurkat 6.3 | Eric Verdin | (Jordan et al., 2003) |
| Jurkat 6.3 NT shRNA | Iván D’Orso | Created in this study |
| Jurkat 6.3 KAP1 shRNA | Iván D’Orso | Created in this study |
| Jurkat 8.4 | Eric Verdin | (Jordan et al., 2003) |
| Jurkat 8.4 NT shRNA | Iván D’Orso | Created in this study |
| Jurkat 8.4 KAP1 shRNA | Iván D’Orso | Created in this study |
| Jurkat 9.2 | Eric Verdin | (Jordan et al., 2003) |
| Jurkat 9.2 NT shRNA | Iván D’Orso | Created in this study |
| Jurkat 9.2 KAP1 shRNA | Iván D’Orso | Created in this study |
| Jurkat 2B2D | Jonathan Karn | (Pearson et al., 2008) |
| Jurkat 2B2D NT shRNA | Iván D’Orso | Created in this study |
| Jurkat 2B2D KAP1 shRNA | Iván D’Orso | Created in this study |
| Jurkat 2B2D (IPTG) NT shRNA | Iván D’Orso | Created in this study |
| Jurkat 2B2D (IPTG) KAP1 shRNA | Iván D’Orso | Created in this study |
| U2OS | ATCC HTB96 | Purchased |
| U2OS NT shRNA | Iván D’Orso | Created in this study |
| U2OS KAP1 shRNA | Iván D’Orso | Created in this study |
| HEK 293T | ATCC CRL-11268 | Purchased |
| HEK 293FT | Thermo Fisher 70007 | Purchased |
| SupT1 | ATCC CRL-1942 | Purchased |
| Jurkat E6.1 | ATCC TIB-152 | Purchased |

**Table S2. shRNA vectors used in this study**

| **Target gene** | **Vector / Restriction sites** | **Primer numbers / sequences (5’-3’) to generate shRNA vectors** |
| --- | --- | --- |
| NT | pLVTHM/ ClaI-MluI | **1342/**CGCGTCCCCCAACAAGATGAAGAGCACCAATTCAAGAGATTGGTGCTCTTCATCTTGTTGTTTTTGGAAAT  **1343/**CGATTTTCAAAAACAACAAGATGAAGAGCACCAATCTCTTGAATTGGTGCTCTTCATCTTGTTGGGGGA |
| KAP1 | pLVTHM / ClaI-MluI | **1338/**CGCGTCCCCCTGAGACCAAACCTGTGCTTATTCAAGAGATAAGCACAGGTTTGGTCTCAGTTTTTGGAAAT  **1339/**CGATTTCCAAAAACTGAGACCAAACCTGTGCTTATCTCTTGAATAAGCACAGGTTTGGTCTCAGGGGGA |
| NELF-E | pLVTHM / ClaI-MluI | **1340/**CGCGTCCCCCTGGATTCCTTGTGCCTCATATTCAAGAGATATGAGGCACAAGGAATCCAGTTTTTGAAAAT  **1341/**CGATTTTCAAAAACTGGATTCCTTGTGCCTCATATCTCTTGAATATGAGGCACAAGGAATCCAGGGGGA |
| NT | pLKO.1 / AgeI-EcoRI | SHC002 (Sigma) |
| KAP1 | pLKO.1 / AgeI-EcoRI | TRCN0000017998 (Sigma)  CCGGCCTGGCTCTGTTCTCTGTCCTCTCGAGAGGACAGAGAACAGAGCCAGGTTTTT |
| NT | pLKO.1-puro-IPTG-3xLacO /  AgeI-EcoRI | SHC332 (Sigma)  CCGGCAACAAGATGAAGAGCACCAACTCGAGTTGGTGCTCTTCATCTGTTGTTTTTG |
| KAP1 | pLKO.1-puro-IPTG-3xLacO /  AgeI-EcoRI | TRCN0000017998 (Sigma)  CCGGCCTGGCTCTGTTCTCTGTCCTCTCGAGAGGACAGAGAACAGAGCCAGGTTTTT |

**Table S3. DNA oligonucleotides used in this study**

*The number of the amplicons used in real-time PCR quantification of the ChIP-enriched DNA represents the midpoint of the two primers respective to the transcription start site (TSS), upstream the TSS (-) and downstream the TSS (+).

**Note that +2627 and +7232 are the same amplicon. +2627 is the position of the amplicon respective to the TSS in the E4 cell-based model and +7232 is the position of the amplicon respective to the TSS in the 10.6 cell-based model and in infection experiments with HIV-1_NL4-3_.

****Note that the +4230 and +9553 are the same amplicon. +4230 is the position of the amplicon respective to the TSS in the E4 cell-based model and +9553 is the position of the amplicon respective to the TSS in the 10.6 cell-based model and in infection experiments with HIV-1_NL4-3_.

| **Amplicon*** | **Primer Number / Sequence (5’-3’)** | **Figure (Assay)** |
| --- | --- | --- |
| -353 | **1093** / AAGGCTACTTCCCTGAT  **1094** / TAGCACCATCCAAAGGTC | 3E, 5F (ChIP) |
| -69 | **1360** / CTTGCTACAAGGGACTT  **1361** / AGGGCTCGCCACTCC | 3E, 5F (ChIP) |
| -37 | **1364** / CTTTCTACAAGGGACTTTCCGCTG  **1365** / CTCCCAGGCTCAGATCTGGTC | 3E, 5F (ChIP) |
| +141  5’-LTR specific | **1111** / GCTTAAGCCTCAATAAAGCTTGCCTTGAG  **1112** / GTCCTGCGTCGAGAGATCTCCTCTG | 3E (ChIP)  4C, 4E, S1D, S1F, S1G (RT-qPCR) |
| +2627 (+7232)** | **1358** / TGAGGGACAATTGGAGAAGTGA  **1359** / TCTGCACCACTCTTCTCTTTGC | 3E, 5F (ChIP)  3C, 4D, 4F, 5E, 7E, S1E, S1F, S1H, S1I, S4B, S4C, S4D, S4E, S4F, S4G (RT-qPCR) |
| +4230  (+9553)***  3’-LTR specific | **1368** / ACAAGAGGAGGAAGAGGTGGGT  **1369** / GCCCTGGTGTGTAGTTCTGCCA | 3E, 5F (ChIP) |
| ACTB | **1256** / GATGATGATATCGCCGCGCT  **1257** / CTTCTCGCGGTTGGCCTTGG | All RT-qPCR experiments |

**Table S4. Antibodies used in this study**

| **Target** | **Company** | **Catalogue Number** | **Assay** |
| --- | --- | --- | --- |
| β-actin (C4) | Santa Cruz Biotechnologies | SC-47778  (1:5000) | Western blot |
| NELF-E (H-140) | Santa Cruz Biotechnologies | SC-32912  (1:2000) | Western blot |
| KAP1 (20C1) | Abcam | AB22553  (1:5000)  (5 μg / 20 million cells) | Western blot  ChIP |
| RNA Pol II (N-20) | Santa Cruz Biotechnologies | SC-899X  (5 μg / 20 million cells) | ChIP |
| Cdk9 (C-20) | Santa Cruz Biotechnologies | SC-484  (5 μg / 20 million cells) | ChIP |
| Normal Mouse IgG | Santa Cruz Biotechnologies | SC-2025  (5 μg / 20 million cells) | ChIP |
| Donkey anti-rabbit IgG-HRP | Santa Cruz Biotechnologies | SC-2313  (1:10.000) | Western blot |
| Goat anti-mouse IgG-HRP | Santa Cruz Biotechnologies | SC-2005  (1:10.000) | Western blot |

**Table S5. Plasmids used in this study**

| **Insert** | **Vector / tag** | **Restriction sites / Reference** |
| --- | --- | --- |
| FFL LUC | pTRIP | SpeI-XhoI / (Schoggins et al., 2011) |
| Tat | pTRIP / STREP | SpeI-XhoI |
| Tat C22G | pTRIP / STREP | SpeI-XhoI |
| KAP1 shRNA | pLVTHM | ClaI-MluI |
| NELF-E shRNA | pLVTHM | ClaI-MluI |
| Non Target (NT) shRNA | pLVTHM | ClaI-MluI |
| KAP1 shRNA | pLKO.1 | AgeI-EcoRI |
| Non Target (NT) shRNA | pLKO.1 | AgeI-EcoRI |
| GAL4 | pcDNA4TO | HindIII-EcoRI |
| GAL4–CycT1 | pcDNA4TO | EcoRI-XhoI |
| GAL4–CDK9 | pcDNA4TO | EcoRI-XhoI |
| GAL4–CDK9 T186A | This study | EcoRI-XhoI |
| HIV LTR – FFL LUC | pcDNA3.1+ | (D'Orso et al., 2012) |
| HIV LTR 5xGal4 – FFL LUC | pcDNA3.1+ | (D'Orso et al., 2012) |
| CMV – RL LUC | pCMV | (D'Orso et al., 2012) |

**Table S6. SDE Parameters Used in the Mathematical Modeling**

Note: The asterisk (*) indicates the only two parameters that change in the four scenarios.

| **Equation** | **Measured parameters** | **Dimension** | **Reference** | **Model Reference** | **Tat+**  **KAP1+** | **Tat-**  **KAP1+** | **Tat+**  **KAP1-** | **Tat-KAP1-** |
| --- | --- | --- | --- | --- | --- | --- | --- | --- |
| dRNA/dt |  |  |  |  |  |  |  |  |
| τ_RNA |  |  |  |  | 0.2071 | 0.2071 | 0.2071 | 0.2071 |
| μ_RNA | 5.00E-002 | transcript/sec | (Tay et al., 2010) | 3 | 3.093 | 3.093 | 3.093 | 3.093 |
| k_Mm | 1.00E+005 | (average # NF-κB in cell) | (Tay et al., 2010) |  | 120 | 120 | 120 | 120 |
| k_synth(h) |  | transcript/sec |  |  | 8.20E-06 | 8.20E-06 | 8.20E-06 | 8.20E-06 |
| k_synth(v) | 0.1 | transcript/sec | (Weinberger et al., 2005) | 6 | 3.93E-04 | 3.93E-04 | 3.93E-04 | 3.93E-04 |
| k_decay | 0.2 | 1/hr | (Reddy and Yin, 1999) | 0.003 | 0.0132 | 0.0132 | 0.0132 | 0.0132 |
| dTat/dt |  |  |  |  |  |  |  |  |
| k_trans | 0.005 - 0.5 | Protein/sec | (Weinberger et al., 2005) | 0.3 - 30 | 0.5412(*) | 0 | 0.5412(*) | 0 |
| μ_Tat | 25-100 | - | (Reddy and Yin, 1999) | 60 | 35.4256 | 35.4256 | 35.4256 | 35.4256 |
| k_MTat |  |  |  |  | 125 | 125 | 125 | 125 |
| d_Tat | 0.154 | Protein/hr | (Reddy and Yin, 1999) | 0.025 | 0.0194 | 0.0194 | 0.0194 | 0.0194 |
| dNF-κB/dt |  |  |  |  |  |  |  |  |
| β | 2e-5 – 0.01 | 1/sec | (Tay et al., 2010) | 0.0012 – 0.6 | 0.5 | 0.5 | 0.5 | 0.5 |
| d_NF-κB | 5.00E-002 | 1/sec | (Tay et al., 2010) | 3 | 3 | 3 | 3 | 3 |
| dKAP1/dt |  |  |  |  |  |  |  |  |
| d_KAP1 |  |  |  |  | 0.006 | 0.006 | 0.006 | 0.006 |
| ρ |  |  |  |  | 0.081(*) | 0.081(*) | 0.001 | 0.001 |
| dTNF/dt |  |  |  |  |  |  |  |  |
| d_TNF | 2.00E-04 | 1/sec | (Tay et al., 2010) | 1.20E-02 | 1.20E-02 | 1.20E-02 | 1.20E-02 | 1.20E-02 |
| dRNAbasal/dt |  |  |  |  |  |  |  |  |
| α | 1.00E-008 | transcript/sec | (Weinberger et al., 2005) | 6.00E-007 | 0.008 | 0.008 | 0.008 | 0.008 |
| d_RNAbasal |  | 1/sec |  |  | 0.00667 | 0.00667 | 0.00667 | 0.00667 |

**Table S7. System of ODEs Describing the Deterministic Approximation of the Mathematical Model**

$$\frac{d\left[ RNA \right]}{dt}=\mu_{RNA}\frac{\left[ NF\kappa B \right]}{k_{Mm}+\left[ NF\kappa B \right]}+\tau_{RNA}\left[ KAP1 \right]\left[ NF\kappa B \right]+k_{synth(h)}\left[ KAP1 \right]\left[ Tat \right]+k_{synth(v)}\left[ Tat \right]-k_{decay}\left[ RNA \right]$$

(1)

$$\frac{d\left[ Tat \right]}{dt}=k_{trans}\left[ RNA \right]+\mu_{Tat}\frac{\left[ Tat \right]}{k_{MTat}+\left[ Tat \right]}-d_{Tat}\left[ Tat \right]$$

(2)

$$\frac{d\left[ NF\kappa B \right]}{dt}=\beta\left[ TNF \right]-d_{NF\kappa B}\left[ NF\kappa B \right]$$

(3)

$$\frac{d\left[ KAP1 \right]}{dt}=\rho-d_{KAP1}\left[ KAP1 \right]$$

(4)

$$\frac{d\left[ TNF \right]}{dt}=-d_{TNF}\left[ TNF \right]$$

(5)

$$\frac{d\left[ RNAbasal \right]}{dt}=\alpha-d_{RNAbasal}\left[ RNAbasal \right]$$

(6)

$$\left[ RNAtot \right]=\left[ RNA \right]+[RNAbasal]$$

(7)
